# The MAPK phosphatase VHP-1 buffers pharynx-to-body proportions against tissue-specific growth imbalance in *C. elegans*

**DOI:** 10.64898/2026.09.28.754458

**Authors:** Ioana Gheorghe, Klement Stojanovski, Delia Bogenstätter, Anna Graf, Dirk Beuchle, Sacha Psalmon, Benjamin Daniel Towbin

## Abstract

Maintaining appropriate organ size ratios in the face of growth fluctuations is critical for the development of a reproducible body plan. Yet the mechanisms involved remain poorly understood. Here, we investigated how pharynx-to-body proportions are maintained in *Caenorhabditis elegans*, combining tissue-specific perturbations, genetic screening, and longitudinal live imaging. A genome-wide RNAi screen revealed that knock-down of the dual-specificity MAPK phosphatase VHP-1 turns animals hypersensitive to inter-tissue growth imbalance caused by pharyngeal or epidermal depletion of the mTORC1 activator RAGA-1 or the ribosomal protein RPL-22. In contrast, *vhp-1* mutants tolerated global *raga-1* loss, indicating a specific requirement for *vhp-1* under tissue growth imbalance. Knock-down of the p38 pathway suppressed the imbalance-specific defects of *vhp-1* mutants. In contrast, JNK knock-down effectively rescued the pleiotropic phenotypes of *vhp-1* mutants but only weakly reduced their sensitivity to RAGA-1 imbalance, indicating that these two stress-MAPK pathways make distinct contributions to the response to growth imbalance. Finally, whole-animal VHP-1 levels increased upon epidermal RAGA-1 depletion, and epidermal VHP-1 depletion did not reproduce the sensitivity caused by global *vhp-1* loss, consistent with a contribution from VHP-1 outside the growth-perturbed epidermis in buffering against local growth imbalance.

**Summary statement:** The MAPK phosphatase VHP-1 is critical for buffering *C. elegans* development against tissue growth imbalance.

*New:* imbalance-vs-slow-growth dissociation, non-autonomy of VHP-1, splitting p38 from JNK functionally, endogenous VHP-1 dynamics.

## Introduction

The size proportions of an animal’s body parts, such as its organs, limbs, and body segments, are strikingly reproducible between individuals (Stern and Emlen, 1999; Vollmer, Casares and Iber, 2017; Boulan and Léopold, 2021; Vea and Shingleton, 2021; Stojanovski *et al*., 2023). For example, in *C. elegans*, pharynx-to-body proportions vary by less than 5% among individuals (Stojanovski *et al*., 2023). This reproducibility is astounding, given that individual organs differ in their intrinsic growth rates and in their sensitivity to nutritional, hormonal, and environmental inputs. All of these differences must be accommodated to faithfully reach correct proportions at adulthood (Andersen, Colombani and Léopold, 2013; Gokhale and Shingleton, 2015; Vea and Shingleton, 2021).

Even pronounced growth defects confined to a single tissue rarely derail the organism as a whole. Instead, tissue-specific perturbations trigger systemic responses that adjust the growth and developmental progression of the whole animal, guiding development toward reproducible outcomes (Parker and Shingleton, 2011; Stojanovski *et al*., 2023). Mechanisms functioning in growth-perturbed tissues are increasingly well described (Colombani, Andersen and Léopold, 2012; Garelli *et al*., 2012; Boone *et al*., 2016; Roselló-Díez *et al*., 2018; Boulan *et al*., 2019; Blanco-Obregon *et al*., 2022). However, how the remaining, unperturbed tissues sense the growth imbalance and appropriately adjust their own growth to preserve size proportions remains much less understood.

Growth-compensatory responses occur across taxa, including in mammalian limb development (Roselló-Díez *et al*., 2018), and mechanistically have arguably been best studied in *Drosophila*. Growth inhibition or damage in a larval imaginal disc slows fly development and delays pupariation (Parker and Shingleton, 2011; Colombani, Andersen and Léopold, 2012; Garelli *et al*., 2012; Gokhale *et al*., 2016), giving the perturbed disc time to complete its growth. Depending on the type of perturbation, this response is triggered by YAP, JNK, or Xrp1 signaling in the damaged disc, converging on secretion of the relaxin-like peptide *Dilp8* (Colombani, Andersen and Léopold, 2012; Garelli *et al*., 2012, 2015; Colombani *et al*., 2015; Vallejo *et al*., 2015; Boulan *et al*., 2019). While *Dilp8* is known to act through the relaxin receptor Lgr3 to suppress biosynthesis of the steroid hormone ecdysone, mechanistic insights into the cellular circuitry that delays growth and development in the receiving, unperturbed tissues remain sparse.

Systemic growth coordination has also been described in *Caenorhabditis elegans*, offering a genetically tractable system in which to address the global response to growth imbalance. Mosaic loss of ribosomal RNA genes arrests growth even of cells that retain functional rRNA genes (Cenik *et al*., 2019), indicating that growth control acts cell non-autonomously also in nematodes. More directly, tissue-specific depletion of the mTORC1 activator RAGA-1 in either the pharynx or the epidermis slows not only the targeted tissue but also the rest of the animal, so that pharynx-to-body proportions are largely preserved despite mTORC1 inhibition being confined to one tissue (Stojanovski *et al*., 2023). This robustness requires the mechano-transducing transcriptional co-activator *yap-1*: animals with reduced *yap-1* function deviate from correct pharynx-to-body proportions and arrest development specifically when challenged with tissue-specific RAGA-1 depletion (Stojanovski *et al*., 2023). This system thus recapitulates, in a tractable genetic setting, the central feature of inter-tissue growth coordination: a systemic response to local growth perturbation, while leaving open how unperturbed tissues mount that response.

Candidate mediators of such a response include the stress-activated MAP kinase (MAPK) pathways JNK and p38, which respond to diverse cellular stresses and regulate physiology, growth, and developmental decisions both cell-autonomously and across tissues (Kim *et al*., 2004; Andrusiak and Jin, 2016; Sanchez *et al*., 2019). In *C. elegans*, the JNK-like MAPK KGB-1 drives cell-non-autonomous stress responses from neurons and other tissues (Liu, Ruediger and Shapira, 2018), while the p38 MAPK PMK-1 couples stress responses to developmental progression (Weaver *et al*., 2020). Sustained activation of these pathways can itself impair development, and their output is tightly constrained by negative regulators, most prominently the dual-specificity MAPK phosphatases (Bermudez, Pagès and Gimond, 2010). In *C. elegans*, the phosphatase VHP-1 antagonizes both JNK and p38 signaling (Kim *et al*., 2004; Mizuno *et al*., 2004): loss of *vhp-1* causes a larval arrest that is suppressed by reducing activity of the JNK/KGB-1 pathway (Mizuno *et al*., 2004), while elevated p38/PMK-1 activity contributes to the developmental delay of *vhp-1* deficient animals (Kim *et al*., 2004; Weaver *et al*., 2020). *vhp-1* thus regulates two separate stress-MAPK branches which, as we show below, show differential involvement in the response to tissue-specific growth imbalance.

In this study, we identify *vhp-1* as a regulator required for robustness to tissue-specific growth imbalance in *C. elegans* in a genome-wide RNAi screen. We find that *vhp-1* becomes critical specifically when growth is imbalanced between tissues and that it is required to preserve both developmental progression and pharynx-to-body size proportions under these conditions. Genetic analysis implicates stress-responsive MAPK signaling, most prominently its p38 branch, in this requirement. Finally, we find that depletion of VHP-1 in the perturbed tissue alone does not recapitulate these phenotypes, suggesting that it has a role outside the growth-perturbed tissue. Together, these findings identify VHP-1-regulated stress-MAPK signaling as a component of the response that allows animals to buffer local growth perturbation and maintain pharynx-to-body proportions.

## Results

### Genome-wide screen identifies synthetic interactors with tissue-specific growth imbalance

To identify molecular regulators of organ growth coordination, we employed a previously established genetic system to create growth imbalance among tissues (Smith *et al*., 2023; Stojanovski *et al*., 2023). An auxin-inducible degradation (AID) tag is used to degrade the mTORC1 activator RAGA-1 specifically in the pharynx by expression of the corresponding E3 ligase TIR1 under the *myo-2* promoter. Despite pharynx-specific knock-down of RAGA-1, pharynx-to-body size proportions remain nearly unchanged, because other tissues are similarly reduced in growth (Stojanovski *et al*., 2023). We reasoned that knockdown of genes required for this systemic growth response should increase the sensitivity of animals to pharyngeal RAGA-1 AID, leading to a disproportionate body plan, developmental defects, or complete larval arrest.

Following this logic, we performed a genome-wide RNAi screen in 384-well plates with quadruplicate repeats for each RNAi clone and condition. Every well was seeded with ∼10 L1 larvae, and the number of progeny per well was counted using automated image analysis four days later. In this assay, a reduction in animal count can result from various defects emerging in the seeded generation (e.g., slow growth, larval arrest, sterility, lethality) or from embryonic lethality of their progeny, and our primary readout does not distinguish between these possibilities. Importantly, however, we applied RNAi in the presence and absence of pharyngeal RAGA-1 AID (***Figure 1****a*). Thus, the screen did not merely identify essential genes, but knockdowns that increased the sensitivity of animals to pharyngeal RAGA-1 depletion.

**Figure 1.**
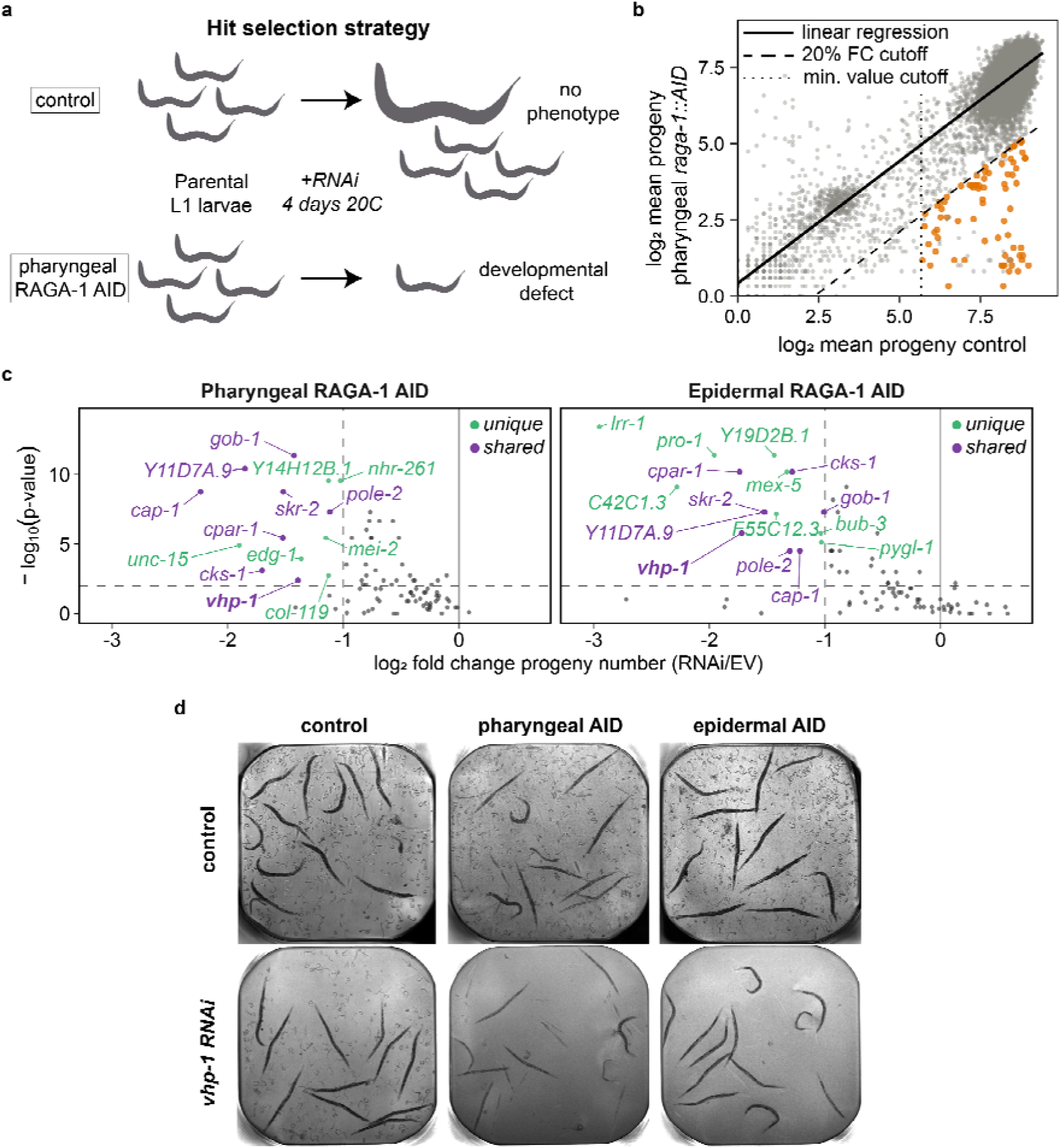
A genome-wide RNAi screen identifies synthetic interactors of tissue-specific RAGA-1 depletion. **a)** Hit selection strategy: L1 larvae of control or pharyngeal RAGA-1 AID animals are fed with RNAi for 4 days at 20°C. For selected candidates, control animals reach adulthood and produce progeny, while pharyngeal RAGA-1 AID animals produce fewer or no progeny due to hypersensitivity to tissue-specific RAGA-1 imbalance. **b)** Genome-wide RNAi screen comparing mean progeny output (log□) for each RNAi clone between animals with pharyngeal RAGA-1 depletion and control animals carrying the raga-1::aid tag, but lacking TIR1 expression. Solid line, robust linear fit across all RNAi clones; dashed line, fold-change threshold (≤0.2 of the value predicted from the control data). Orange points indicate hits selected for further study based on the following criteria: FDR-adjusted p < 0.005, fold change ≤ 0.2, and control progeny ≥ 50. Error! Reference source not found. provides the complete list of primary hits, including gene names, WormBase IDs, fold changes, and p-values. **c)** Combined volcano plots showing results from the validation screen of initial hits in pharyngeal and epidermal RAGA-1 AID backgrounds. Both backgrounds were compared with control animals carrying raga-1::aid but lacking TIR1 expression. Each point represents one RNAi clone; the log fold change in progeny output relative to the value predicted from the matched control data, normalized using the empty-vector (EV) relationship between strains, is plotted against −log (p-value; one-sided binomial test). Dashed lines indicate the significance thresholds (p < 0.01 and fold change ≤ 0.5). Green: hits unique to the indicated tissue-specific background; purple: hits significant in both backgrounds; grey: not significant. The complete list of validated unique and shared hits, including log fold changes and p-values, is provided in Error! Reference source not found.. **d)** Representative images of animals with unperturbed RAGA-1 (control) or with RAGA-1 AID in indicated tissues following control or vhp-1(RNAi) treatment.

In the absence of RNAi, pharyngeal RAGA-1 AID was permissive for development but reduced the animal count 2.8-fold (***Figure 1****b*, **Error! Reference source not found.**a). To identify screen hits, we therefore scored knockdowns that caused at least a 5-fold bigger reduction in animal count under pharyngeal RAGA-1 AID compared to control conditions 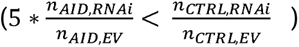. To exclude genes with strong pleiotropic phenotypes, we additionally excluded RNAi treatments that reduced the animal count to less than 50 in the control conditions from further analysis.

We re-tested the 73 strongest candidate genes (*Error! Reference source not found.*) from the genome-wide screen and additionally assessed a potential synthetic interaction of each RNAi with epidermal, instead of pharyngeal RAGA-1 AID. We thereby distinguished candidates specifically sensitive to pharyngeal RAGA-1 impairment from those that were also affected by reduced RAGA-1 activity in another tissue, making them more likely to be involved in a general response to growth imbalance rather than pharynx-specific mechanisms. This secondary screen validated a total of 8 RNAi clones that showed increased sensitivity to both pharyngeal and epidermal RAGA-1 AID (minimal fold change: 2-fold, *p<0.01*, ***Figure 1****c*, Error! Reference source not found.).

Among the validated candidates, we focused on *vhp-1,* a dual-specificity phosphatase (DUSP) of MAP kinases orthologous to human DUSP8/16 (Mizuno *et al*., 2004; Bye-A-Jee *et al*., 2020). VHP-1 dephosphorylates the JNK and p38 MAPKs *in vitro* (Mizuno *et al*., 2004), and its loss causes larval arrest that is genetically suppressed by reducing activity of the KGB-1/JNK pathway (Mizuno *et al*., 2004). VHP-1 also negatively regulates the PMK-1/p38 pathway, as *vhp-1* depletion increases PMK-1 phosphorylation and the resulting developmental defects are partially suppressed by mutation of *pmk-1* (Kim *et al*., 2004; Weaver *et al*., 2020; Yuan *et al*., 2023). This established role of *vhp-1* as a negative regulator of stress-responsive MAPK signaling, together with the sensitivity of *vhp-1(RNAi)* animals for both types of tissue-specific perturbations in our validation assay (***Figure 1****d*), made it an attractive candidate for a signaling regulator involved in organ growth coordination.

### *vhp-1* is required for robustness of growth and development to growth imbalance between tissues

To validate and further characterize the role of *vhp-1* in the response to tissue-specific RAGA-1 imbalance, we used longitudinal live imaging (Turek, Besseling and Bringmann, 2015; Gritti *et al*., 2016; Stojanovski, Großhans and Towbin, 2022). Individual eggs were placed in agarose-based microchambers and imaged every 10-12 minutes from hatching throughout larval development. For imaging, animals expressed two fluorescent reporters: a green fluorescent protein expressed in the pharyngeal muscle (*myo-2p::gfp*) and a ubiquitously expressed red fluorescent protein (*eft-3p::mScarlet*). Automated image analysis using a custom-made pipeline (Materials and Methods) enabled extraction of body and pharynx size, growth rates, and individual growth trajectories throughout larval development. Growth plateaus associated with molting further provided estimates of larval stage durations and developmental arrest (***Figure 2****a*).

**Figure 2.**
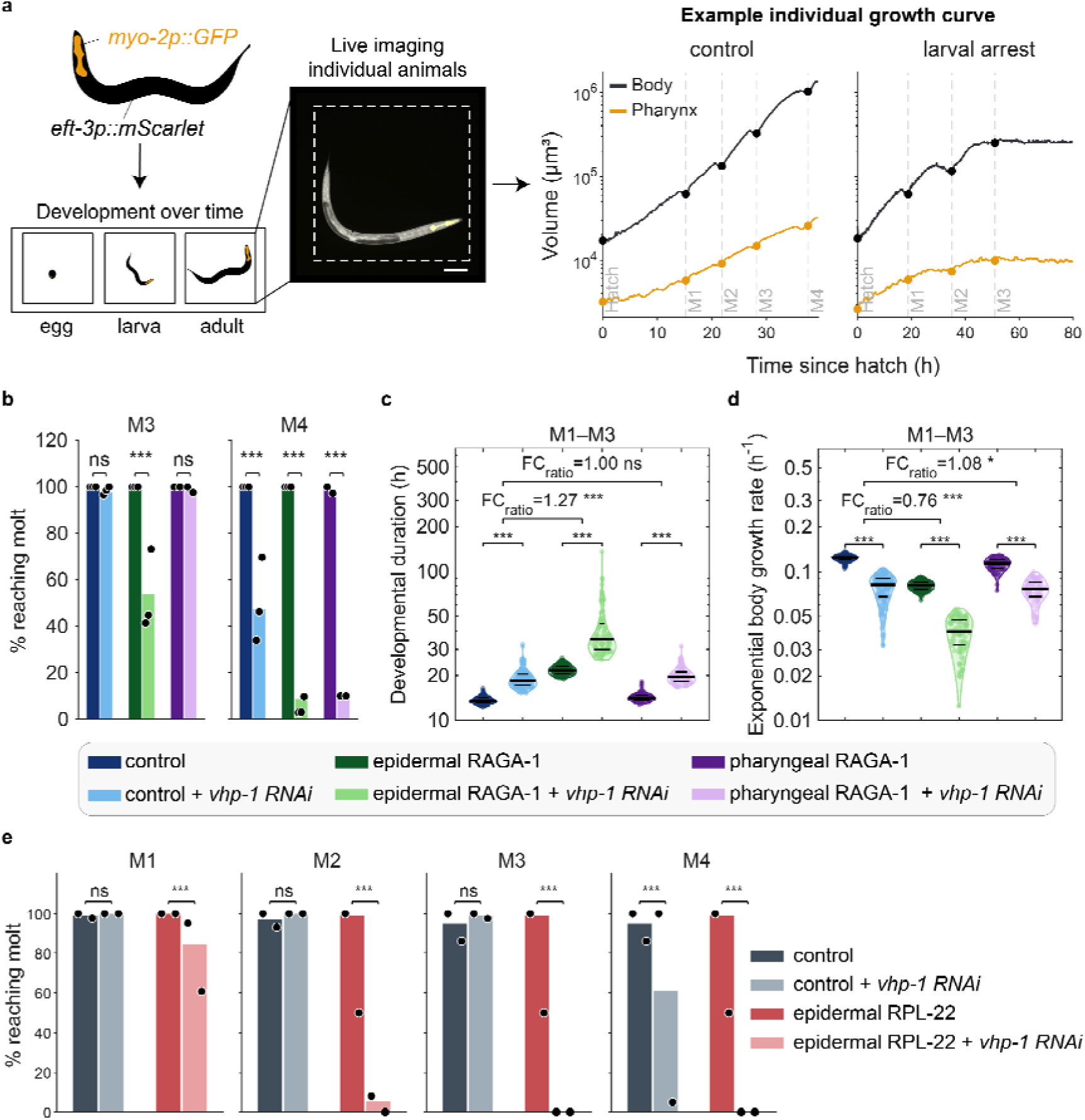
vhp-1 is required for robustness of developmental speed and larval stage progression to tissue-specific growth inhibition. **a)** Experimental approach for longitudinal imaging of individual animals expressing pharyngeal myo-2p::GFP and ubiquitous eft-3p::mScarlet. Eggs were loaded into 600 × 600 × 20 µm agarose microchambers and imaged every 10– 12 min. Representative body and pharynx volume trajectories are shown for a control animal and an animal undergoing larval arrest. Dashed vertical lines indicate larval molts M1–M4. Scale bar, 100 µm. **b)** Percentage of animals reaching M3 and M4 in indicated conditions. Black circles indicate day-to-day repeats. Bars: mean of day-to-day repeats. **c)** Developmental duration and exponential body growth rate between M1 and M3. Violin plots show the distributions of individual animals; horizontal lines indicate the first quartile, median, and third quartile. Colors indicate conditions as described in the gray box below the graph. Statistics: pairwise comparisons: two-sided Wilcoxon rank-sum tests; Comparison of effect size between pairs: two-way ANOVA on log-transformed values followed by linear contrasts. FC ratios represent the vhp-1(RNAi) vs. control fold change in each tissue-specific RAGA-1 condition divided by the corresponding fold change in the control RAGA-1 condition. Missing and non-positive values were excluded, and extreme outliers were removed from display on the log scale using a three-IQR threshold. **e)** Percentage of animals reaching M1–M4 under control conditions or epidermal RPL-22 depletion at 10 µM auxin, with or without vhp-1(RNAi). Within each condition, control and vhp-1(RNAi) were compared using two-sided Fisher’s exact tests. Sample sizes, pairwise fold changes, and exact p-values for b–e are provided in Error! Reference source not found.. ns, p ≥ 0.05; *p < 0.05; **p < 0.01; ***p < 0.001.

To enhance *vhp-1* knockdown efficiency during early L1 development, we applied RNAi already from the L4 stage of the parental generation. Under these conditions, ∼50% of the animals did not complete the fourth molt (M4), even without additional perturbation by RAGA-1 AID, consistent with the larval arrest reported for genetic *vhp-1* null mutants (Mizuno *et al*., 2004). However, this phenotype was significantly enhanced by tissue-specific RAGA-1 AID: depletion of RAGA-1 in either the pharynx or the epidermis reduced the fraction of animals completing M4 to below 10%, and epidermal RAGA-1 AID additionally impaired completion of the preceding molt (M3) (***Figure 2****b*). Tissue-specific depletion of RAGA-1 in an otherwise wild-type background did not impact developmental success, showing that *vhp-1* is required for developmental robustness to these growth perturbations.

Even in larval stages preceding arrest, animals with combined epidermal RAGA-1 depletion and *vhp-1(RNAi)* had a longer delay in development between M1 and M3 upon epidermal RAGA-1 AID (60.2% increase in double depleted animals vs. 36.8% in *vhp-1(RNAi)* alone, *p = 2.745e-23,* ***Figure 2****c*), and a corresponding slow-down of the exponential volume growth rate (***Figure 2****d*, p = 3.208e-23). By contrast, *vhp-1(RNAi)* combined with pharyngeal RAGA-1 depletion caused substantial larval arrest, but animals that completed development were not measurably delayed (*p = 0.902*), suggesting both parallels and differences in the role of *vhp-1* in responding to pharyngeal and epidermal RAGA-1 perturbations.

To test if the sensitivity of *vhp-1(RNAi)* animals to RAGA-1 AID represented a specific interaction with the mTORC1 pathway or also occurred for other mechanisms of tissue-growth imbalance, we epidermally depleted the ribosomal protein RPL-22 by AID. When titrating the auxin to a concentration that reduced the growth rate to a similar degree as epidermal RAGA-1 AID (***Supplementary Figure S2*** *a, b*), nearly 100% of the individuals successfully completed development in otherwise unperturbed conditions. However, less than 5% of *vhp-1(RNAi)* treated animals reached M2, and no animal reached M3 when exposed to the same dose of auxin (***Figure 2****e*).

Together, these data confirm the sensitivity of *vhp-1(RNAi)* animals to pharyngeal RAGA-1 AID and extend these observations to epidermal depletion of RAGA-1 and RPL-22. This broad sensitivity of *vhp-1(RNAi)* animals to different growth perturbations suggests a sensitivity to tissue growth imbalance rather than a specific interaction of *vhp-1* with *raga-1* or its involvement in the development of a specific tissue.

### *vhp-1* is critical for maintaining pharynx-to-body proportions upon epidermal RAGA-1 inhibition

To test whether *vhp-1(RNAi)* also affected the robustness of body-plan proportions to tissue-specific RAGA-1 inhibition, we measured pharynx and body length, width, and volume at each molt from individual growth trajectories and quantified the deviation in pharynx-to-body proportion with and without tissue-specific RAGA-1 AID. Importantly, since the pharynx and body grow allometrically, they naturally change in proportion during development (Stojanovski *et al*., 2023). Treatments that alter the body size therefore also alter adult pharynx-to-body proportions, even if scaling to body size is unperturbed. To correct these effects, we compared the observed pharynx size in each individual to the size predicted for its body size by a log-linear model fitted to measurements from a corresponding control condition (***Figure 3****a*).

**Figure 3.**
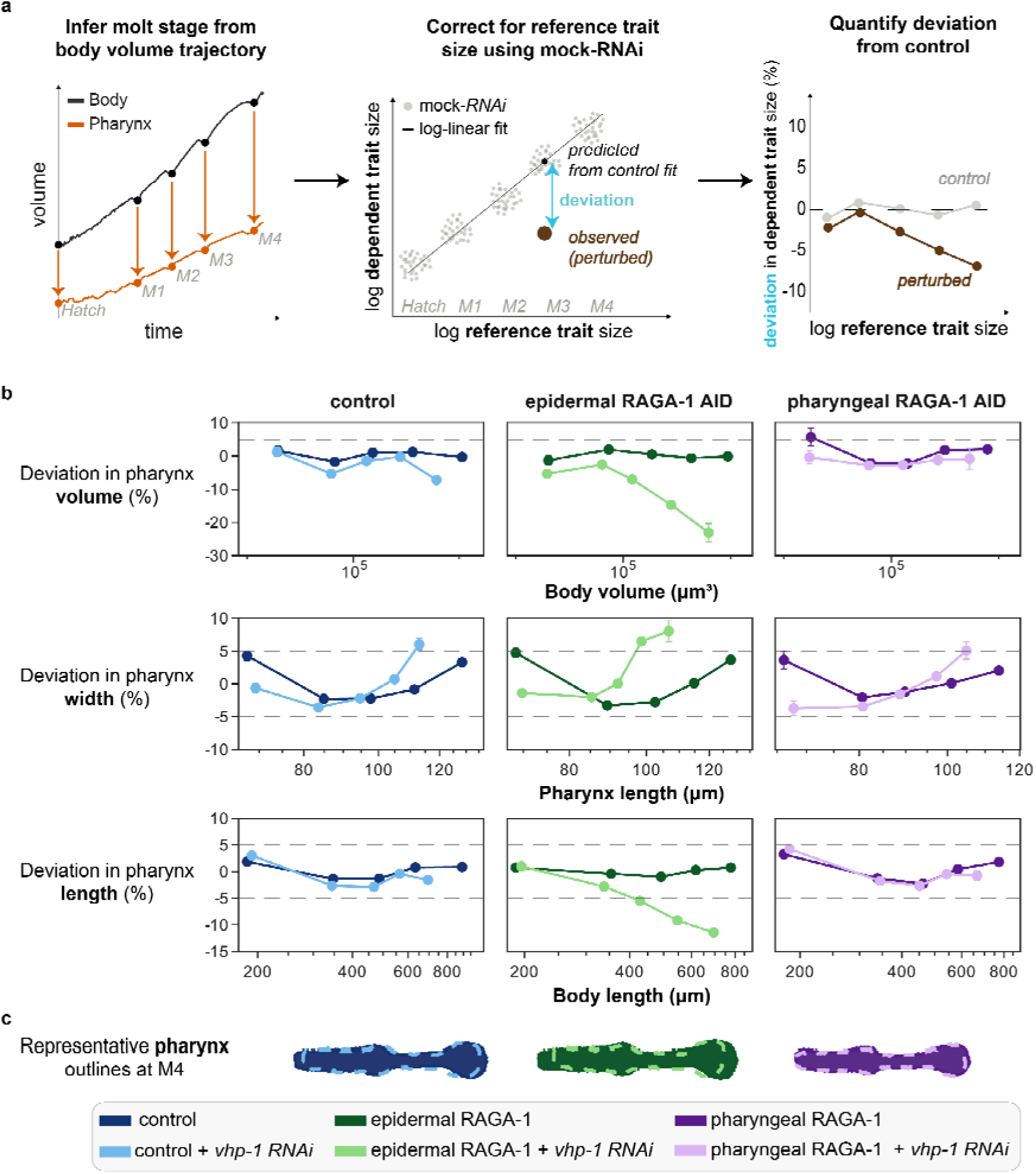
vhp-1 is required for robustness of pharynx-to-body proportions to epidermal RAGA-1 AID. **a)** Schematic of the analysis workflow. Body volume size trajectories were used to identify molt stages. To account for overall size changes, a log–log model relating each dependent pharyngeal trait (e.g., pharynx length) to the corresponding reference trait (e.g., body length) was fitted for mock-RNAi control animals. For each individual, the percentage deviation of the observed pharyngeal measurement from the value predicted by the control-fitted model was then calculated. **b)** Mean percentage deviation in pharynx volume (top), width (middle), and length (bottom) in indicated conditions. Pharynx volume and length deviations are plotted against body volume and body length, respectively, whereas pharynx width deviation is plotted against pharynx length. Separate control log– log models were fitted for each experimental set. Each point represents the mean deviation across all individuals measured at one ecdysis. Error bars: SEM across individuals. Where no error bars are visible, they are smaller than the marker. Dashed horizontal lines indicate ±5% deviation from the control prediction. Per-stage sample sizes and exact p-values are provided in Error! Reference source not found.. **c)** Representative pharynx outlines at M4 from animals matched for body length. For each genetic background, the reference body length was defined as the mean M4 body length of the corresponding vhp-1(RNAi) group, and representative animals were selected to match this reference body length as closely as possible. Colors indicate conditions as defined in the legend.

Accounting for altered body size, *vhp-1(RNAi)* only weakly affected pharynx-to-body length proportions in wild-type animals or in animals with pharyngeal RAGA-1 depletion. By contrast, *vhp-1(RNAi)* reduced pharynx-to-body proportions upon epidermal RAGA-1 depletion (***Figure 3****b*) by more than 10 %. Similarly, deviations in pharynx length-to-width proportions and in body-to-pharynx volume proportions were specifically enhanced when *vhp-1(RNAi)* was combined with epidermal RAGA-1 AID. The combined perturbation of *vhp-1* and epidermal RAGA-1 rendered pharynxes more stunted (increased width-to-length ratio) and decreased the overall pharynx-to-body volume proportions (***Figure 3****c*).

Together, these data show that *vhp-1* is required to buffer pharynx-to-body proportions against growth perturbation by epidermis-specific RAGA-1 depletion.

### Tissue-specific, but not global *raga-1* impairment enhances *vhp-1-*dependent developmental defects

To further investigate the role of *vhp-1* in the robustness to epidermal growth imbalance, we examined the partial hypomorphic allele *vhp-1(sa366)*. This allele introduces a premature stop codon that truncates the C-terminal region of VHP-1, while leaving the catalytic domain intact (***Figure 4****a*). Unlike *vhp-1* null mutants, which arrest during larval development (Mizuno *et al*., 2004), *vhp-1(sa366)* animals remain viable in an otherwise wild-type background (Choy and Thomas, 1999).

**Figure 4.**
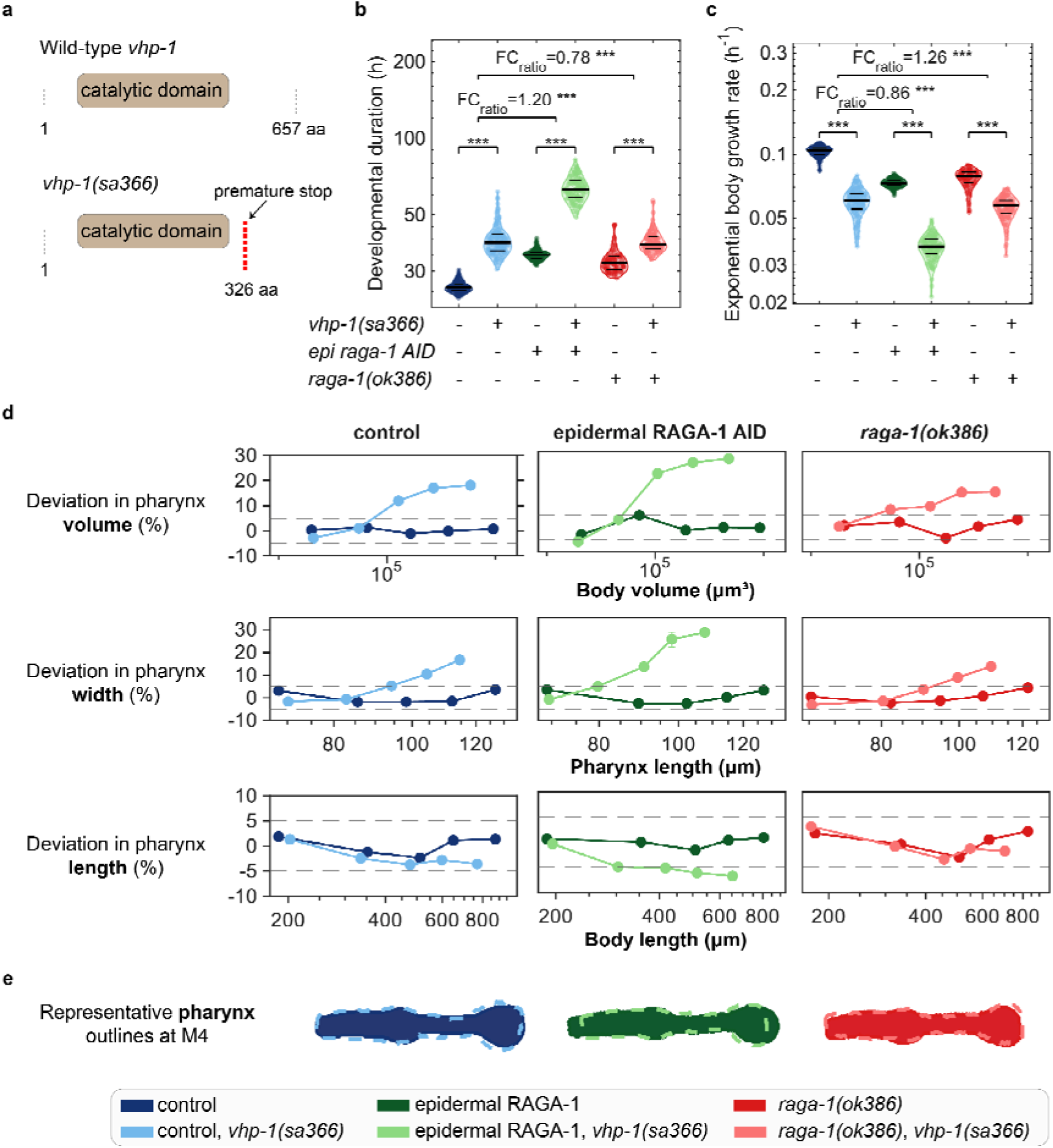
vhp-1 mutant is hypersensitive to epidermal RAGA-1 AID, but not to ubiquitous raga-1 loss. **a)** Schematic representation of predicted protein encoded by wild-type vhp-1 and vhp-1(sa366) mutant. The sa366 mutation introduces a premature stop codon. **b, c)** Developmental duration and exponential growth rate from M1–M4 of indicated strains and conditions. Violin plots show the distributions of individual animals; horizontal lines indicate the first quartile, median and third quartile. Fold-change ratios (FC ratio) and p-values for the indicated contrasts are shown above the plots. Statistical tests as in Fig. 2c. ***p < 0.001. **d)** Percentage deviation in pharynx volume (top), width (middle) and length (bottom) from the relationship predicted by the corresponding control-fitted log–log model. Dark lines represent unperturbed vhp-1, light lines vhp-1(sa366). Each point represents the mean deviation at one ecdysis, and error bars indicate SEM across individuals. Where no error bars are visible, they are smaller than the marker. Dashed horizontal lines: ±5% deviation from the regression model to the control. **e)** Representative pharynx outlines at M4 from animals matched for body length. For each genetic background, the reference body length was defined as the mean M4 body length of the corresponding vhp-1(sa366) group; within each vhp-1 genotype, the animal with body length closest to this reference value was selected. Colors indicate conditions as defined in the legend. Sample sizes, pairwise fold changes, and exact p-values for b–d are provided in Error! Reference source not found.. ***p < 0.001.

Live imaging showed that, similar to *vhp-1(RNAi)*, the *vhp-1(sa366)* allele increased the time to complete development from M1 to M4 by 49.8% (***Figure 4****b*) and correspondingly reduced the average exponential growth rate over this period (***Figure 4****c*). Like for *vhp-1(RNAi)* animals, the growth rate of *vhp-1(sa366)* mutants was synergistically reduced upon epidermal RAGA-1 AID (1.519-fold in unperturbed versus 1.808-fold upon epidermal RAGA-1 AID; *p = 4.402e-90*), although the mutant allele did not cause larval arrest (***Supplementary Figure S3****a)*. Critically, this enhanced delay was specific to tissue-specific RAGA-1 loss: it did not occur with global loss of *raga-1* by the ubiquitous deletion allele *raga-1(ok386)*, even though the global *raga-1* loss by itself slowed growth to a similar extent as epidermis-specific depletion (***Figure 4****b, c*). This specificity of the phenotype to tissue-specific depletion points to a role of *vhp-1* in the response to tissue-growth imbalance, rather than a general genetic interaction with mTORC1 signaling across all tissues.

Measurements of pharynx and body size confirmed the role of *vhp-1* in buffering length proportions against epidermis-specific RAGA-1 depletion and additionally revealed a critical role in the maintenance of pharynx shape. As for *vhp-1(RNAi)*, mutation of *vhp-1(sa366)* specifically reduced the pharynx length when combined with epidermal RAGA-1 AID, but not as strongly in wild-type animals, or in ubiquitous *raga-1(ok386)* mutants (***Figure 4****d*). Compared to RNAi, the increase in pharyngeal width was more pronounced in *vhp-1(sa366)* mutants (***Figure 4****e*), such that the pharyngeal volume proportions were increased rather than decreased in the *vhp-1(sa366)* mutant, despite its shortened pharyngeal length. Consistent with RNAi results, however, the effects of the *vhp-1(sa366)* mutation were synergistically enhanced by epidermal RAGA-1 AID, but not by ubiquitous mutation of *raga-1*.

In summary, we consistently find synthetic interactions between epidermal RAGA-1 AID and *vhp-1* for RNAi and the *sa366* allele. The different effects of the mutant and RNAi on pharynx width may reflect differences in residual catalytic activity, or a defect specific to the truncated protein of the *sa366* allele. Importantly, we show that the sensitivity of *vhp-1* mutants is restricted to tissue-specific imbalance of RAGA-1 and does not occur upon global *raga-1* loss.

### Upstream JNK and p38 regulators suppress *vhp-1* growth delay, but only p38 suppresses the scaling defect upon epidermal growth inhibition

VHP-1 is known to dephosphorylate both JNK and p38 MAPKs *in vitro*, and the larval arrest of *vhp-1* null mutants is suppressed by depleting or by mutating components of the JNK/p38 pathways (Kim *et al*., 2004; Mizuno *et al*., 2004; Weaver *et al*., 2020; Yuan *et al*., 2023). We therefore asked whether the sensitivity of *vhp-1(sa366)* mutants to growth imbalance was likewise suppressed by knock-down of JNK or p38 pathways. To identify such suppressors, we depleted upstream regulators and downstream effectors of both pathways by RNAi and tested whether any of these knockdowns enhanced the growth of epidermal RAGA-1 AID; *vhp-1(sa366)* double perturbed animals.

Equivalent to the genome-wide screen described above (***Figure 1***), animals were grown in 384-well plates in quadruplicate repeats per RNAi clone, and the number of animals was scored after four days of incubation (***Figure 5****a*). This screen identified 10 strong suppressors that increased the animal count after 4 days by more than 4-fold (*p< 5e-6*) in epidermal RAGA-1 AID; *vhp-1(sa366)* double perturbed animals (***Figure 5****b*). Suppressors included components of both the JNK and p38 pathways, as well as the transcription factor *daf-16/FOXO*. While *daf-16* functions as an effector of the insulin-like signaling pathway, it has also been reported to interact genetically with stress-associated MAPK signaling (Kondo *et al*., 2005; Oh *et al*., 2005; Twumasi-Boateng *et al*., 2012).

**Figure 5.**
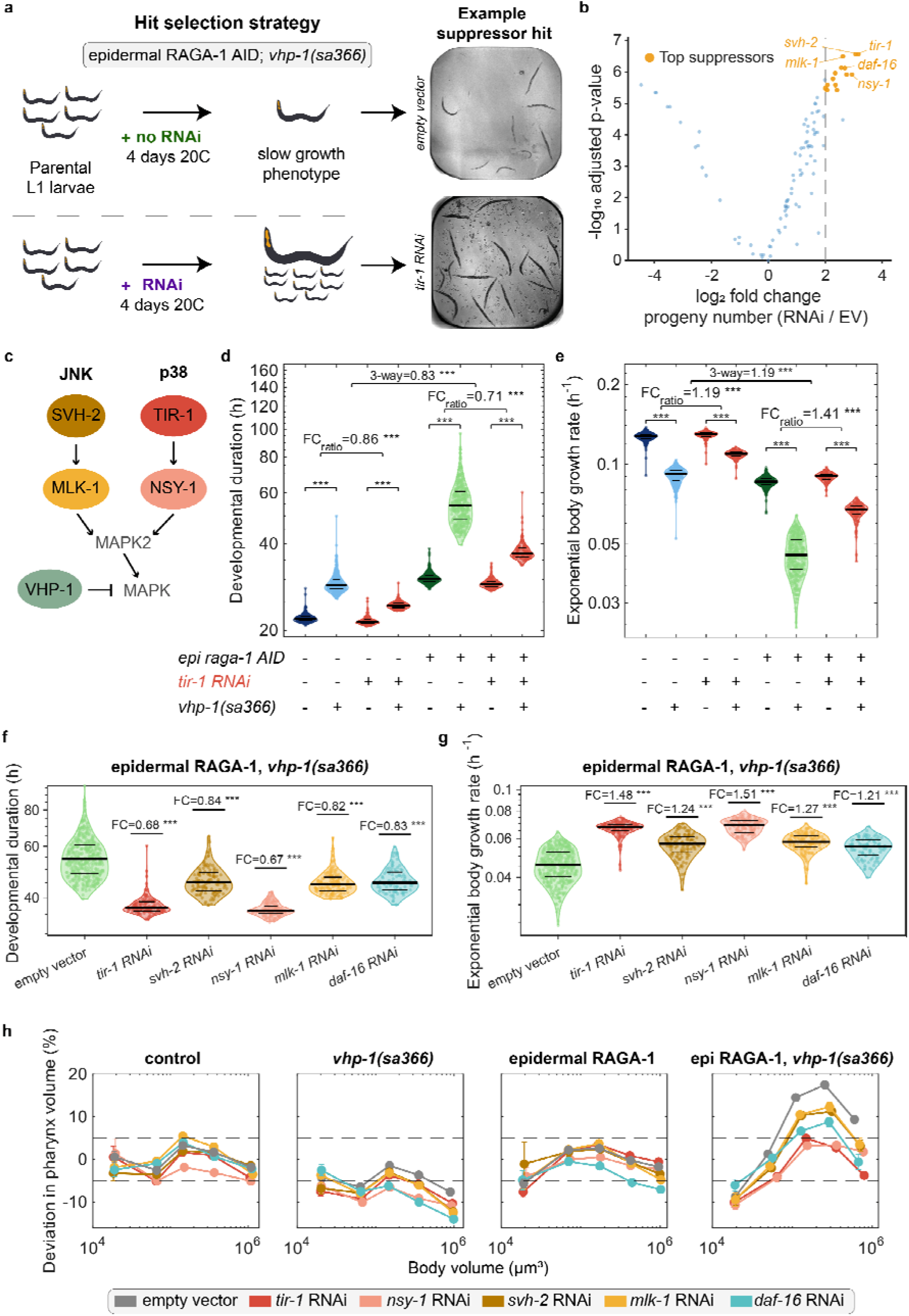
JNK and p38 pathway inhibition suppress vhp-1-associated growth delay and pharynx–body proportion defects with stronger suppression by p38 inhibition. **a)** Suppressor-screen strategy. Parental L1 larvae carrying epidermal RAGA-1 AID and vhp-1(sa366) were cultured for 4 days at 20 °C in 384-well plates. Suppressors were called as RNAi that increased progeny production compared to mock RNAi. **b)** RNAi suppressor screen in epidermal RAGA-1 AID; vhp-1(sa366) animals. Each point represents one RNAi clone; the log2 fold change in progeny number relative to empty vector (EV) RNAi is plotted against the −log_10_ adjusted p-value. Orange points indicate the strongest suppressors, defined by a greater than two-fold increase in progeny number and an adjusted p-value below 5*10^−6^. Dashed line: two-fold-change threshold. **c)** Simplified representation of the JNK and p38 MAPK pathways showing the suppressors selected for longitudinal analysis. SVH-2 and MLK-1 act upstream of JNK signaling, whereas TIR-1 and NSY-1 act upstream of p38 signaling. **d, e)** Developmental duration and exponential growth rate from M1–M4. Violin plots show the distributions of individual animals; horizontal lines indicate the first quartile, median, and third quartile. Fold-change ratios (FC ratio) and p-values for the indicated contrasts are shown above the plots. “3-way” denotes the interaction among epidermal RAGA-1 depletion, vhp-1(sa366) and tir-1(RNAi). Statistics as in Figure 2. **f, g)** Direct comparison of M1–M4 developmental duration and exponential growth rate, respectively, following inhibition of the indicated JNK and p38 pathway components. Plot elements are as in d, e; FC denotes median fold change relative to EV. Two-sided Wilcoxon rank-sum tests versus EV. **h)** Percentage deviation in pharynx volume from the relationship predicted by the control-fitted regression model, plotted against body volume, in control, vhp-1(sa366), epidermal RAGA-1 AID, and epidermal RAGA-1 AID; vhp-1(sa366) animals for indicated conditions and RNAi treatments. Each point represents the mean deviation at one ecdysis. Error bars SEM across individuals. Where no error bar is visible, it is smaller than the marker. Dashed horizontal lines: ±5% deviation from the control prediction. Sample sizes and exact statistical results for d–h, together with the corresponding analyses in **Supplementary Figures S4–6**, are provided in Error! Reference source not found.. ***p < 0.001.

To confirm these results, we conducted growth assays in microchambers for five of the strongest suppressors. For the JNK pathway, we tested the putative receptor *svh-2* and the MAP3K *mlk-1*. For the p38 pathway, we assessed the upstream regulator *tir-1* and the MAP3K *nsy-1*. Additionally, we included *daf-16* as a potential downstream effector (***Figure 5****c*).

Depletion of either JNK- or p38-pathway components suppressed the growth delay of *vhp-1(sa366)* mutants (***Figure 5****d, e*, ***Error! Reference source not found.****a-c,* ***Supplementary Figure S5****a-c**Error! Reference source not found.***) for strains with and without epidermal RAGA-1 AID. Importantly, however, the suppression was significantly more pronounced for the doubly perturbed strain (*p= 6.575e-128*), suggesting that reduction of p38/JNK signaling also suppressed the synergistic interaction of *vhp-1* and epidermal RAGA-1 imbalance, rather than merely a general defect of the *vhp-1* mutant alone.

While RNAi of either branch, p38 and JNK, yielded significant suppression, the effect size was much bigger for RNAi against the p38 branch (*tir-1* and *nsy-1*) than for JNK (*mlk-1* and *svh-2*) (***Figure 5****f,g*). This stronger suppression suggests a more important role of p38 signaling in robustness to epidermal RAGA-1 imbalance compared to JNK, although we cannot entirely exclude that this difference is also driven by technical effects, such as RNAi efficiency, or genetic redundancy. Similar to JNK, *daf-16(RNAi)* suppressed the general *vhp-1(sa366)* delay and had a weaker effect on the epidermal RAGA-1 AID-specific genetic interaction (***Error! Reference source not found.****d,* ***Supplementary Figure S5****d*).

The role of p38 signaling in robustness to growth imbalance was also apparent from pharynx-to-body size measurements. RNAi of neither p38 nor JNK components altered pharynx-to-body volume proportions in wild-type animals or in epidermal RAGA-1 AID animals. Both moderately reduced these proportions in the *vhp-1(sa366)* mutant alone. However, the deviation in volume proportion caused by combined epidermal RAGA-1 AID and *vhp-1(sa366)* mutation was nearly completely rescued by RNAi of p38 components (***Figure 5****h*). The deviation in pharyngeal length and width was also substantially rescued, although neither measure fully returned to control values and slight deviations remained (***Error! Reference source not found.6****a, b*). In contrast, RNAi of JNK components (*svh-2*; *mlk-1*) did not fully restore volume proportions (***Figure 5****h*). RNAi of *daf-16* produced an intermediate, partial rescue of pharynx-to-body proportions (***Figure 5****h*), suggesting that other effectors may also be involved. The more pronounced suppression of pharynx-to-body scaling defects by knock-down of p38-pathway components compared to the JNK pathway suggests a stronger contribution of p38 signaling to the growth-imbalance related phenotypes of *vhp-1* mutants.

### VHP-1 levels oscillate across the molt cycle and increase upon epidermis-specific RAGA-1 AID

*vhp-1* transcription is induced by JNK activity following mechanical trauma in *C. elegans* (Egge *et al*., 2021), and the mammalian homolog DUSP16 is transcriptionally induced by stress-MAPK signaling (Han, Kim and Heasley, 2002; Lee *et al*., 2010). As a stress-MAPK phosphatase whose own expression is coupled to MAPK activity, VHP-1 could itself be dynamically regulated during development. We therefore asked whether VHP-1 protein levels change in response to tissue-specific growth imbalance.

To measure VHP-1 expression, we endogenously tagged VHP-1 with the green fluorescent protein mStayGold (mSG). Consistent with previous transcriptional reporters (Mizuno *et al*., 2004), VHP-1:mSG was broadly expressed across tissues (***Figure 6****a*, **Supplementary Figures S7***a*). The tagged allele was functional, as the mSG tagged strain showed no slowdown of growth upon epidermal RAGA-1 AID (**Supplementary Figure S7***b*). For reliable quantification in microchambers, we imaged the tagged strain in parallel with a matched untagged control to determine and subtract bacterial and worm autofluorescence (**Supplementary Figure S7***c*). Additionally, all strains carried a single-copy *emr-1::mCherry* nuclear marker (Morales-Martínez, Dobrzynska and Askjaer, 2015) to control for changes in detection efficiency during development.

**Figure 6.**
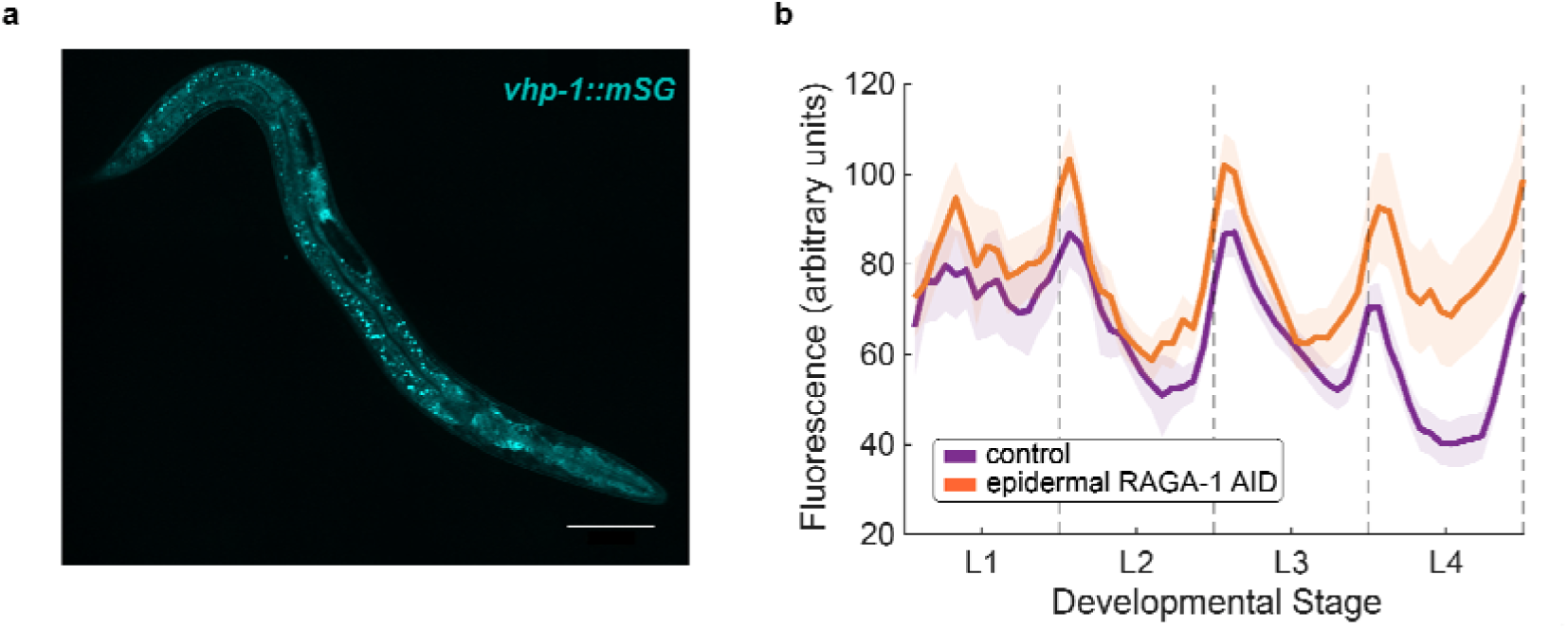
VHP-1:mStayGold expression oscillates during larval development and increases upon epidermal RAGA-1 AID. **a)** Representative maximum-intensity projection of an L2 animal carrying endogenously tagged vhp-1::mStayGold. Scale bar, 50 µm. **b**) Background-subtracted VHP-1:mStayGold fluorescence concentration across larval development in control animals (N =4, n = 108) and animals with epidermal RAGA-1 AID (N =4, n = 96). Lines show mean fluorescence over the top 5 brightest slices of the z-stack; shaded regions indicate 95% confidence intervals. Dashed vertical lines: molt boundaries. Individuals were rescaled and aligned to the beginning and end of each molt before averaging.

Volumetric confocal time-lapse imaging revealed that VHP-1:mSG levels oscillated, peaking shortly after each molt (***Figure 6****b*), consistent with available transcriptomic data (***Error! Reference source not found.****d*) (Meeuse *et al*., 2023). This oscillation did not simply reflect concentration changes caused by the volumetric growth pause at each molt, as the control protein EMR-1:mCherry oscillated far more weakly (**Supplementary Figure S7***e*). VHP-1 abundance therefore oscillates in a molt-coupled manner during larval development. Quantitative comparison of the oscillatory signal showed that VHP-1:mSG was mildly but reproducibly increased upon epidermal RAGA-1 depletion (***Figure 6****b*), consistent with VHP-1 responding to epidermal growth imbalance.

We were unable to corroborate these dynamics with direct measurements of JNK/p38 MAPK phosphorylation: experimental noise, together with the transient and potentially oscillating nature of the signal, precluded reliable quantification of p38 and JNK phosphorylation by immunoblot. VHP-1 levels therefore provide our most reliable *in vivo* readout of stress-MAPK regulation, albeit an indirect one. The increase under growth imbalance is consistent with VHP-1 tracking stress-MAPK activity in a negative-feedback relationship, although we cannot exclude contributions from other inputs to *vhp-1* expression.

Notably, our assay quantified VHP-1:mSG across the whole animal rather than in individual tissues. However, because the epidermis constitutes only a small fraction of the sampled volume (**Supplementary Figure S7***c*), the increase in total VHP-1:mSG upon epidermal RAGA-1 depletion is consistent with VHP-1 upregulation extending beyond the perturbed epidermis. Although our whole-animal measurements cannot quantitatively assign this increase to individual tissues, spatial inspection of the whole-animal signal did not indicate hotspots of upregulation, consistent with a broadly distributed rather than tissue-restricted induction.

### VHP-1 is not specifically required in the epidermis for the growth-imbalance response

To test whether *vhp-1* is required specifically in the epidermis or systemically across tissues, we used AID to deplete VHP-1 in the epidermis with the same *col-10p::TIR1* driver used for epidermal RAGA-1 AID, and measured growth in microchambers (***Figure 7****a*). Applied alone, epidermal VHP-1 depletion reduced growth rate and extended developmental duration relative to controls, similar to ubiquitous *vhp-1* mutation or RNAi. Co-depletion of RAGA-1 in the epidermis reduced this growth rate further (***Figure 7****b*). Critically, however, unlike for the whole-body depletion of *vhp-1*, epidermal VHP-1 and RAGA-1 AID had a near-additive, rather than synergistic effect: both perturbations independently slowed growth, but the VHP-1 AID effect was of similar magnitude with and without epidermal RAGA-1 depletion (FC of 1.140 vs. 1.163) with only a minimal, albeit statistically significant (*p = 5.912e-08*), interaction between the two treatments. This near independence of the growth effects of RAGA-1 and VHP-1 AID suggests that *vhp-1* function in the epidermis alone does not account for its role in robustness to epidermal growth imbalance.

**Figure 7.**
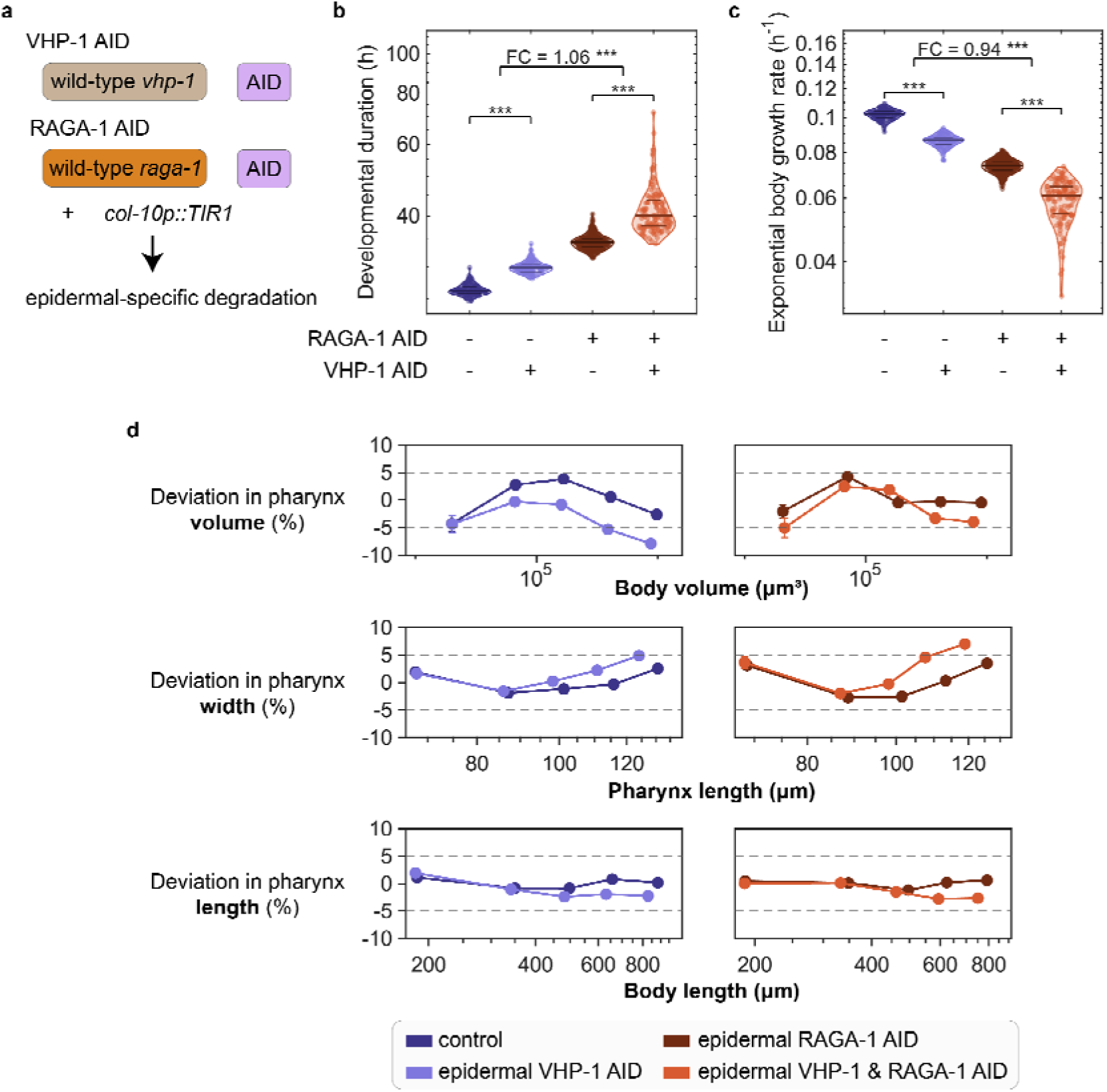
Depletion of VHP-1 specifically in the epidermis does not cause hypersensitivity to epidermal RAGA-1 AID. **a)** Strategy for epidermis-specific co-depletion of VHP-1 and/or RAGA-1 by AID using col-10p::TIR1. **b, c)** Developmental duration and exponential growth rate from M1–M4. Violin plots show the distributions of individual animals; horizontal lines indicate the first quartile, median and third quartile. Fold changes and interaction-test p values are indicated. Statistics as in Figure 2, ***p < 0.001. **d)** Percentage deviation in pharynx volume, length and width from the relationship predicted by the corresponding log–log model. Pharynx volume and length deviations are plotted relative to body volume and body length, respectively, whereas pharynx width deviation is plotted relative to pharynx length. Unlike ubiquitous vhp-1 RNAi or mutation, epidermis-specific VHP-1 depletion altered pharynx-to-body proportions similarly in animals with and without epidermal RAGA-1 depletion. Dashed horizontal lines indicate ±5% deviation from the control prediction. Per-stage sample sizes and exact p-values are provided in Error! Reference source not found.. ***p < 0.001.

This interpretation was confirmed by direct measurement of pharynx-to-body size proportions upon epidermal VHP-1 AID. Epidermis-specific depletion of VHP-1 slightly reduced the pharynx-to-body proportions relative to control (*p < 0.001*), confirming that the VHP-1 depletion was functionally effective. However, unlike what we observed for ubiquitous mutation or depletion of *vhp-1* (***Figure 3****b*, ***Figure 4****d*), this effect was not enhanced or even weaker when RAGA-1 was simultaneously epidermally depleted by AID. Thus, unlike the ubiquitous *vhp-1(sa366)* mutation, epidermal VHP-1 depletion did not interact synergistically with epidermal RAGA-1 imbalance.

These results suggest that *vhp-1* function in response to organ growth imbalance is not primarily in the epidermis, i.e. not in the tissue targeted by the growth inhibition. Instead, these data are consistent with a contribution of VHP-1 outside the growth-perturbed epidermis to developmental robustness.

## Discussion

Animals must coordinate the growth of their organs to reach reproducible body proportions, even when individual tissues are perturbed by injury, mutation, or local variation in nutrient and growth signaling (Andersen, Colombani and Léopold, 2013; Gokhale and Shingleton, 2015; Vea and Shingleton, 2021). Such coordination must distinguish imbalance in growth between tissues from a global, coordinated reduction in the growth rate (Parker and Shingleton, 2011; Stojanovski *et al*., 2023) and must adjust the growth of different tissues appropriately.

Here, we identify the dual-specificity MAP kinase phosphatase VHP-1 as a regulator of this response in *Caenorhabditis elegans*. Reducing VHP-1 activity, by RNAi or mutation, sensitized animals to tissue-specific growth inhibition caused by epidermal or pharyngeal depletion of the mTORC1 activator RAGA-1, or of the ribosomal protein RPL-22, causing developmental delay, a disproportionate pharynx-to-body scaling, or larval arrest. Critically, this sensitivity was specific to local perturbation of *raga-1*, whereas global mutation of *raga-1* did not synergistically enhance *vhp-1* defects in growth or body plan proportions. VHP-1 therefore acts to buffer development against imbalance between tissues, rather than as a general facilitator of growth.

VHP-1 dephosphorylates both the JNK and p38 stress-activated MAP kinases (Kim *et al*., 2004; Mizuno *et al*., 2004), and consistent with this dual activity, reducing signaling through either branch suppressed the general growth impairment and developmental delay of *vhp-1* mutants. The two branches were, however, not equivalent: knockdown of p38-pathway components rescued the defects arising from combined mutation of *vhp-1* and epidermal RAGA-1 depletion near completely, whereas only very weak suppression was observed by JNK-pathway knock-down. Since both kinases are VHP-1 substrates and both contribute to the generic consequences of VHP-1 loss, this dissociation associates the proportion-restoring function primarily to the p38 branch, rather than to generic stress-MAPK hyperactivation. We additionally recovered DAF-16/FOXO as a *vhp-1* partial suppressor of the body plan scaling defects. As DAF-16 has been reported to act with stress-associated MAPKs (Oh *et al*., 2005; Twumasi-Boateng *et al*., 2012; Liu, Ruediger and Shapira, 2018) in addition to its role downstream of insulin-like signaling (Lin *et al*., 1997; Ogg *et al*., 1997; Murphy and Hu, 2018), these data hint that the *vhp-1* mediated response converges in part on FOXO-dependent transcription.

Where does VHP-1 act to maintain balanced tissue growth? An obvious possibility was that it functions cell-autonomously within the growth-impaired epidermis to restrain a local stress response, but our data argue against this model. Epidermis-specific VHP-1 depletion was functionally effective in slowing down growth on its own. Yet, unlike ubiquitous VHP-1 depletion, these phenotypes combined near-additively with epidermal RAGA-1 depletion and did not enhance the pharyngeal scaling defect. These results suggest that VHP-1 also functions in one or more tissues other than the perturbed epidermis, consistent with the reported cell-non-autonomous action of both stress-MAPK branches in *C. elegans* (Liu, Ruediger and Shapira, 2018; Foster *et al*., 2020; Weaver *et al*., 2020; Yuan *et al*., 2023). Imaging of endogenously tagged VHP-1 reinforced this conclusion: VHP-1 was broadly expressed and whole-animal VHP-1 levels increased upon epidermal RAGA-1 depletion. Because the epidermis comprises only a small fraction of the sampled volume, this increase is consistent with induction extending beyond the perturbed tissue. Together, these genetic and imaging data support a contribution of VHP-1 outside the perturbed tissue to the response to growth imbalance.

These observations suggest a parsimonious model. Stress-MAPK signaling is known to induce *vhp-1* transcription: JNK drives *vhp-1* expression after mechanical trauma in *C. elegans* (Egge *et al*., 2021), and the mammalian orthologue DUSP16/MKP-7 is likewise stress-induced (Matsuguchi *et al*., 2001; Niedzielska *et al*., 2014; Egge *et al*., 2021), placing VHP-1 in a negative-feedback loop that restrains the kinases that activate it. We propose that localized growth impairment may elevate stress-MAPK signaling, mounting a proportion-restoring growth response and inducing VHP-1 to keep this response within bounds. Partial loss of VHP-1 would then weaken this feedback arm, leaving stress-MAPK signaling inadequately restrained and the response mis-scaled.

Inter-organ growth coordination has been documented across animals, and our findings both parallel and extend this work. In fetal mice, mosaic cell-cycle arrest in left-limb chondrocytes preserves limb symmetry through local compensatory proliferation coupled to a systemic reduction in the growth of unperturbed tissues (Roselló-Díez *et al*., 2018). This local-plus-systemic logic parallels the contribution of VHP-1 outside the growth-perturbed tissue suggested by our data. In *Drosophila*, growth-impaired imaginal discs secrete the relaxin-like peptide Dilp8, which acts through ecdysone modulation to slow distant tissues and preserve proportions (Colombani, Andersen and Léopold, 2012; Garelli *et al*., 2012, 2015; Colombani *et al*., 2015; Vallejo *et al*., 2015). Stress-MAPK signaling has been similarly implicated here: over-expression of the JNK phosphatase Puckered (Martín-Blanco *et al*., 1998), a functional counterpart of VHP-1, in ribosomal-protein-depleted tissue suppresses Dilp8 induction (Colombani, Andersen and Léopold, 2012), though a distinct paradigm engages Dilp8 also via Xrp1 largely independently of JNK (Boulan *et al*., 2019). Strikingly, Puckered and VHP-1 act in a mirror-image logic: in flies, added phosphatase activity dampens the coordinating signal in the source tissue, whereas our data in worms are consistent with an important contribution of VHP-1 outside the growth-perturbed tissue. Stress-MAPK phosphatases thus appear to be a recurring node in growth coordination, even if the tissue in which they act, and the branch they restrain, may differ between species.

Several aspects of our model remain untested. We have not measured p38 or JNK activation directly: bulk approaches are confounded by the pathway’s molt-coupled oscillation in conjunction with the developmental heterogeneity of epidermally RAGA-1 depleted *vhp-1* mutants, and available antibodies could not resolve MAPK activity spatially. Fluorescent MAPK biosensors (Regot *et al*., 2014; de la Cova *et al*., 2017), measured by the longitudinal imaging approach we use here, may offer a route to map where and when each branch is engaged in future work. Assigning the tissue in which VHP-1 acts will likewise require tissue-specific rescue experiments, although interpretability could be challenged by potential overexpression phenotypes (Egge *et al*., 2021). Finally, we have not identified the signal that reports epidermal growth impairment to other tissues. In *Drosophila* this role is filled by secreted Dilp8, but no equivalent is known in worms, and the initiating cue, be it humoral, metabolic, or mechanical, remains the central open question raised by this work.

In summary, we identify VHP-1 as a regulator of developmental robustness to tissue-specific growth imbalance in *C. elegans*. VHP-1 becomes particularly important under tissue-specific growth perturbation but not under comparably growth-slowing global *raga-1* loss, and our data support a contribution from VHP-1 outside the growth-perturbed tissue. Genetic suppression further points to a preferential involvement of the p38 branch of stress-MAPK signaling in this response. Together with Dilp8 signaling in *Drosophila* (Colombani, Andersen and Léopold, 2012; Garelli *et al*., 2012, 2015; Colombani *et al*., 2015; Vallejo *et al*., 2015; Boulan *et al*., 2019) and local-plus-systemic compensation in the mouse limb (Roselló-Díez *et al*., 2018), our findings support the view that coordinating growth to preserve size proportions is a conserved feature of animal development, with stress-associated MAPK signaling as a recurring node within it.

## Materials and Methods

### C. elegans strains

The *vhp-1(aBT17)* [*vhp-1::AID::HiBiT*] and *vhp-1(syb9598)* [*vhp-1::mStayGold*] alleles were generated in this study by CRISPR/Cas9 genome editing. All other strains were constructed by genetic crossing using the previously described alleles and transgenes: *raga-1(wbm40)* (Smith *et al*., 2023), *raga-1(ok386)* (The C. elegans Deletion Mutant Consortium, 2012), *vhp-1(sa366)* (Choy and Thomas, 1999), *ieSi60* (Zhang *et al*., 2015), *reSi1* (Ashley *et al*., 2021), *bqSi577* (Toker *et al*., 2022), *wbmIs88* (Silva-García *et al*., 2019), and *bqSi225* (Morales-Martínez, Dobrzynska and Askjaer, 2015).

wBT160 *raga-1(wbm40) [raga-1::AID::EmGFP] II; bqSi577 [myo-2p::GFP + unc-119(+)] IV; wbmIs88 [eft-3p::3xFLAG::dpy-10 crRNA::SL2::wrmScarlet::unc-54 3’ UTR] V:8645000*.

wBT182 *raga-1(wbm40) [raga-1::AID::EmGFP] II. (3.41) ieSi60[myo-2p::TIR1::mRuby::unc-54 3’UTR+Cbr.unc-119(+)] II (1.73); ?unc-119(ed3)? III; bqSi577 [myo-2p::GFP + unc-119(+)] IV; wbmIs88 [eft-3p::3xFLAG::dpy-10 crRNA::SL2::wrmScarlet::unc-54 3’ UTR] V:8645000*.

wBT186 *reSi1 [col-10p::TIR1::F2A::mTagBFP2::NLS::AID::tbb-2 3’UTR] (I:-5.32); raga-1(wbm40) [raga-1::AID::EmGFP] II; bqSi577 [myo-2p::GFP + unc-119(+)] IV; wbmIs88 [eft-3p::3xFLAG::dpy-10 crRNA::SL2::wrmScarlet::unc-54 3’ UTR] V:8645000*.

wBT344 *raga-1(wbm40) [raga-1::AID::EmGFP] II; bqSi577 [myo-2p::GFP + unc-119(+)] IV; wbmIs88 [eft-3p::3xFLAG::dpy-10 crRNA::SL2::wrmScarlet::unc-54 3’ UTR] V:8645000; vhp-1 (sa366) II*.

wBT338 *reSi1 [col-10p::TIR1::F2A::mTagBFP2::NLS::AID::tbb-2 3’UTR] (I:-5.32); raga-1(wbm40) [raga-1::AID::EmGFP] II; bqSi577 [myo-2p::GFP + unc-119(+)] IV; wbmIs88 [eft-3p::3xFLAG::dpy-10 crRNA::SL2::wrmScarlet::unc-54 3’ UTR] V:8645000; vhp-1 (sa366) II*.

wBT394 *raga-1(ok386) II; vhp-1 (sa366) II; bqSi577 [myo-2p::GFP + unc-119(+)] IV; wbmIs88 [eft-3p::3xFLAG::dpy-10 crRNA::SL2::wrmScarlet::unc-54 3’ UTR] V:8645000*.

wBT395 *raga-1(ok386) II; bqSi577 [myo-2p::GFP + unc-119(+)] IV; wbmIs88 [eft-3p::3xFLAG::dpy-10 crRNA::SL2::wrmScarlet::unc-54 3’ UTR] V:8645000*.

wBT356 *rpl-22(ohm9)[rpl-22::AID::3xFLAG] II; bqSi577 [myo-2p::GFP + unc-119(+)] IV; wbmIs88 [eft-3p::3xFLAG::dpy-10 crRNA::SL2::wrmScarlet::unc-54 3’ UTR] V:8645000*.

wBT369 *rpl-22(ohm9)[rpl-22::AID::3xFLAG] II; bqSi577 [myo-2p::GFP + unc-119(+)] IV; wbmIs88 [eft-3p::3xFLAG::dpy-10 crRNA::SL2::wrmScarlet::unc-54 3’ UTR] V:8645000; reSi1 [col-10p::TIR1::F2A::mTagBFP2::NLS::AID::tbb-2 3’UTR] (I:-5.32)*.

wBT470 *raga-1(wbm40) [raga-1::AID::EmGFP*] II; bqSi225[pBN34(unc-119(+) emr-1p::emr-1::mCherry)] IV; vhp-1(syb9598) II*.

wBT471 *reSi1 [col-10p::TIR1::F2A::mTagBFP2::NLS::AID::tbb-2 3’UTR] (I:-5.32); raga-1(wbm40) [raga-1::AID::EmGFP*] II; bqSi225[pBN34(unc-119(+)emr-1p::emr-1::mCherry)] IV; vhp-1(syb9598) II*.

wBT447 *raga-1(wbm40) [raga-1::AID::EmGFP*] II, bqSi225[pBN34(unc-119(+) emr-1p::emr-1::mCherry)] IV*.

wBT449 *reSi1 [col-10p::TIR1::F2A::mTagBFP2::NLS::AID::tbb-2 3’UTR] (I:-5.32), raga-1(wbm40) [raga-1::AID::EmGFP*] II, bqSi225[pBN34(unc-119(+) emr-1p::emr-1::mCherry)] IV*.

wBT439 *reSi1 [col-10p::TIR1::F2A::mTagBFP2::NLS::AID::tbb-2 3’UTR] (I:-5.32), bqSi577 [myo-2p::GFP + unc-119(+)] IV; wbmIs88 [eft-3p::3xFLAG::dpy-10 crRNA::SL2::wrmScarlet::unc-54 3’ UTR] V:8645000*.

wBT489 *vhp-1(aBT17) [vhp-1::AID::hibit] II; raga-1(wbm40) [raga-1::AID::EmGFP] II; wbmIs88 [eft-3p::3xFLAG::dpy-10 crRNA::SL2::wrmScarlet::unc-54 3’ UTR] V:8645000; bqSi577 [myo-2p::GFP + unc-119(+)] IV*.

wBT487 *vhp-1(aBT17) [vhp-1::AID::hibit] II; wbmIs88 [eft-3p::3xFLAG::dpy-10 crRNA::SL2::wrmScarlet::unc-54 3’ UTR] V:8645000; bqSi577 [myo-2p::GFP + unc-119(+)] IV*.

wBT488 *vhp-1(aBT17) [vhp-1::AID::hibit] II; reSi1 [col-10p::TIR1::F2A::mTagBFP2::NLS::AID::tbb-2 3’UTR] (I:-5.32); wbmIs88 [eft-3p::3xFLAG::dpy-10 crRNA::SL2::wrmScarlet::unc-54 3’ UTR] V:8645000; bqSi577 [myo-2p::GFP + unc-119(+)] IV*.

wBT490 *vhp-1(aBT17) [vhp-1::AID::hibit] II; raga-1(wbm40) [raga-1::AID::EmGFP] II; reSi1 [col-10p::TIR1::F2A::mTagBFP2::NLS::AID::tbb-2 3’UTR] (I:-5.32); wbmIs88 [eft-3p::3xFLAG::dpy-10 crRNA::SL2::wrmScarlet::unc-54 3’ UTR] V:8645000; bqSi577 [myo-2p::GFP + unc-119(+)] IV*.

### RNAi in liquid

RNAi clones from the Ahringer RNAi library (Kamath *et al*., 2003) were replicated from frozen stocks onto LB agar supplemented with Ampicillin (100 µg/mL ) in OmniTrays (Thermo Scientific Nunc, catalog no. 734-0490) and incubated overnight at 37 °C. Subsequently, each colony was used to inoculate 300 µL LB medium in 96-well polypropylene plates (Thermo Scientific Abgene, catalog no. AB-1127), sealed with gas-permeable membranes and incubated overnight at 37 °C with orbital shaking at 180 rpm. Double-stranded RNA expression was induced by supplementation of isopropyl β-D-1-thiogalactopyranoside (IPTG) to a final concentration of 4 mM to the saturated overnight culture and incubation for an additional 4 h at 37 °C with shaking at 180 rpm. Cultures were pelleted by centrifugation at 3’000 rpm for 15 min at room temperature. Supernatants were removed by inverting the plates and tapping onto absorbent paper to remove residual medium. Bacterial pellet was resuspended in 100 µl S basal supplemented with 100 µg/ml carbenicillin, 5 µg/ml cholesterol, and 4 mM IPTG.

Gravid adults were subjected to alkaline hypochlorite treatment, and the recovered eggs were passed through a 40-µm cell strainer, followed by overnight incubation at 25 °C in cholesterol-free S basal with continuous rotation to obtain arrested L1 larvae. 10-µl of L1 larvae were dispensed at 1 larva/µl in S basal supplemented with 100 µg/ml carbenicillin, 5 µg/ml cholesterol, 4 mM IPTG, 2mM indole-3-acetic acid (IAA), and 0.01% Triton X-100 to each well of a flat bottom 384-well assay plate. 10 µl of bacterial suspension was added to each well, splitting each well of a 96well plate into four quadrants of a 384well assay plate. Assay plates were covered with lids and placed in sealed, humidified containers and incubated for 4 days at 20 °C.

Prior to imaging, 80 µL of 12.5 mM levamisole prepared in S basal was added to each well. Plates were imaged using a Nikon Eclipse Ti2 inverted epifluorescence microscope equipped with a 4x objective (NA = 0.25) and a Hamamatsu ORCA-Flash4.0 sCMOS camera. Images were acquired in three channels: brightfield, a green fluorescence channel for detection of pharyngeal *myo-2p::GFP*, and a red fluorescence channel for detection of the body-wide mScarlet signal.

Screening was conducted in daily batches of 13 RNAi library source plates per strain. Each batch included one control plate carrying the empty vector (EV), and positive controls (*ama-1* and *mex-3* RNAi).

### Image and data analysis of RNAi in liquid

For the genome-wide screen, the pharynx label from *myo-2p*:GFP was used to count the number of animals. To pharynxes, a segmentation model was trained using Ilastik (Berg *et al*., 2019), and a binary mask was created from the probability in MATLAB using global thresholding. Adult pharynxes were excluded by overlap with separately segmented adult body signal from the red channel and a size threshold (min. 130, max 1,000 pixels perimeter). The same procedure was applied for screen validation, except that binary pharynx segmentation was directly create in Ilastik and adult pharynxes were not excluded. Prior to validation, individual clones were obtained from the genome-wide library and sequence verified.

For each RNAi clone, the mean progeny count was calculated separately for WBT160 and WBT182 across the four technical replicate wells per strain. Mean progeny counts were transformed as log (*x* + 1). To model the expected relationship between progeny production in the two strains, robust linear regression was performed using the rlm function from the MASS R package. The log -transformed WBT182 progeny count was specified as the response variable, and the corresponding log -transformed WBT160 progeny count was specified as the explanatory variable. For each RNAi clone, a one-sided binomial test was used to assess whether progeny counts in WBT182 were significantly lower than predicted by the regression model by testing all 16 pairwise comparisons per RNAi clone of the quadruplicate repeats of the two strains. The resulting *P* values were adjusted for multiple testing using the Benjamini–Hochberg false-discovery-rate procedure.

An effect-size measure was calculated as the ratio of the observed mean progeny count in WBT182 to the corresponding value predicted by the robust regression model:

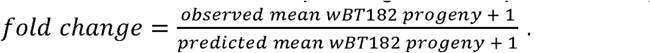

RNAi clones were classified as primary hits if they met all three of the following criteria: a Benjamini–Hochberg-adjusted *P* value of less than 0.005, a fold change of 0.2 or less, and a mean progeny count of at least 50 in control conditions (WBT160).

### Statistics analysis validation screen

Strains WBT182 and WBT186 were each compared separately with the control strain WBT160. Statistical analysis was identical to the genome-wide screen, except that instead of comparing individual clones to a regression to the full data, a regression was fit to empty vector control, forcing to go through the origin. RNAi clones were classified as validated hits if they met all three of the following criteria: a one-sided binomial *P* value of less than 0.01, a fold change of 0.5 or less, and a mean pharynx count of at least 50 in wBT160.

### *vhp-1* suppressor identification

The suppressor screen was performed using strain WBT338 in the presence of IAA, with empty-vector (EV) bacteria serving as the negative control. Animals were incubated with the RNAi bacteria for 5 days at 20 °C before immobilization and imaging. Otherwise, experimental and image analysis procedures were as described for the validation screen. The identities of the 10 highest-ranked suppressor clones were subsequently verified by Sanger sequencing. The remaining clones were not sequence-verified.

Progeny number was normalized to the adult count. Wells in which no adults were detected were excluded from the analysis. Statistically significant suppressors were determined by a two-sided Wilcoxon rank-sum test with adjustment for multiple testing using the Benjamini– Hochberg false-discovery-rate procedure (*P* < 5 × 10^−^□) and a greater than 4-fold increase in progeny per adult relative to the empty-vector control.

### Assembly of agarose microchambers for long-term live imaging

For all experiments, the parental generation was grown on standard plates at 25 °C and embryonic progeny was used to load microchambers. For RNAi experiments, eggs were transferred from NGM plates seeded with OP50-1 to RNAi plates containing 4 mM IPTG and 100 µg/mL carbenicillin seeded with HTT115 carrying the relevant RNAi clone at L4 stage and eggs of the next generation were collected for loading into the microchambers.

Agarose microchambers were prepared as described in Stojanovski et al., 2023 (Stojanovski *et al*., 2023). Microchamber dimensions were 600 x 600 x 20 µm, except for *vhp-1:mSG* expression measurement, for which 300 x 300 x 20 µm was used.

Imaging of growth measurements was carried out on a Nikon Ti2 epifluorescence microscope equipped with a 10x objective (NA = 0.45) and a Hamamatsu ORCA-Flash4 sCMOS camera (pixel size of 6.5 µm). A feedback-controlled IceCube temperature-regulation system (Life Imaging Services) maintained the microscope at 25 °C throughout the experiments. GFP and mScarlet were excited using independently controlled 470-nm and 575-nm LEDs, respectively, from a Spectra X light source (Lumencor). TTL triggering was used to minimize delays when switching between excitation wavelengths. The exposure time was 10 ms per channel, and each two-color acquisition was completed within 30 ms. Software-based autofocus was performed in NIS-Elements (Nikon) every 10–12 min using low-intensity 575-nm illumination. For *vhp-1:mStayGold* measurement, a Nikon Ti2 microscope equipped with a CSU-W1 spinning-disk confocal unit, a 40x objective (NA = 1.3), and a dual-Kinetix22 camera configuration (Photometrics) camera using 2 x 2 camera binning was used. Fluorescence z-stacks with 41 optical sections planes acquired at 1-µm intervals were acquired at hourly intervals for fluorescence intensity measurements. Additionally, single plane acquisitions in wide-field mode were acquired at 20-min time intervals to monitor growth. Samples were maintained at 25 °C throughout imaging using an incubator encasing the microscope (Life Imaging Services).

### Image analysis

Images acquired from animals in standard 600 x 600 x 20-µm microchambers were analyzed using a custom-made pipeline. For link to code and segmentation model weights, see the Code availability section.

For each animal and time point, body and pharyngeal morphology was segmented in 2D from *eft-3p::mScarlet* and *myo-2p::GFP* channels using a deep learning segmentation model trained using manually annotated images. Both masks were computationally straightened prior to morphological analysis of length, area, and width. Width was the average number of pixels in dorso-ventral direction. Volume was estimated from the straightened masks assuming rotational-symmetry.

Automated quality-control using an XGBoost classifier trained with manually annotated input images was used to exclude poor quality images and to distinguish hatched worms from eggs. Animals where the bacterial food supply was depleted before completion of development were manually identified and were excluded from the analysis. Molting time points were detected using a trained 1D neural network applied to the body volume trajectories and manually inspected and corrected if necessary using a custom-made graphical user interface.

### Calculation of developmental duration and growth rates

Developmental durations were calculated for each animal as the sum of the durations of the larval stages contained within the specified developmental interval. Animals for which one or more stage durations within the analyzed interval were missing were excluded from that calculation.

Exponential body growth rates were calculated for each animal between the beginning and end of the specified developmental interval as: 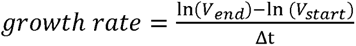, where V and V_end_ are the estimated body volumes at the two developmental landmarks and Δt is the total duration of the respective larval stages in hours. Animals with missing or non-positive volume measurements, missing stage durations or a non-positive total duration were excluded from the corresponding calculation.

Extreme outliers in developmental duration and growth rate were identified separately within each condition using the log-transformed values. Values below Q1 − 3 x IQR or above Q3 + 3 x IQR were excluded from plotting and statistical analysis.

To quantify deviations of one morphological trait relative to another, a scaling relationship was fitted separately for each experimental comparison using the corresponding control animals. For example, the relationship between pharyngeal volume, P, and body volume, B, was fitted by linear regression in log–log space: ln(*P*) = *a* + *b* * ln(*B*). The model was fitted using measurements from all control animals and molt timings included in the analysis. For each animal and its respective molts, the percentage deviation from the value predicted by the control relationship was calculated as: 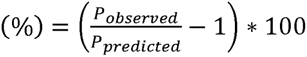.

### Analysis of *vhp-1:mStayGold* expression

Brightfield images and confocal fluorescence z-stacks were processed independently, with separate deep-learning segmentation models trained on manually annotated images acquired in the 300 x 300 x 20-µm microchambers. For the brightfield images, total body masks were generated using a model trained specifically on brightfield images from these experiments. The growth trajectories were visually inspected and used to manually annotate hatching and completion of the four larval molts for each animal.

Confocal z-stacks were segmented using a separate deep-learning model trained on the *vhp-1::mStayGold* fluorescence signal. Segmented area and mean fluorescence intensity were extracted for each z-plane using custom MATLAB scripts. For each animal and time point, the five z-planes with the highest mean *vhp-1::mStayGold* fluorescence intensity were selected, and fluorescence measurements from these planes were averaged. Five planes were used consistently because they represented approximately 20% of the worm-containing z-planes across the experimental time course. Analyses using alternative numbers and proportions of z-planes produced consistent results. Where the *emr-1::mCherry* channel was analyzed, fluorescence was measured at the same z-positions selected using the *vhp-1::mStayGold* signal.

Fluorescence trajectories were aligned to developmental stage using hatching and molt times annotated from brightfield images. For each animal, measurements within each larval stage were linearly interpolated onto 15 evenly spaced developmental-time points. Reporter-specific *vhp-1::mStayGold* fluorescence was calculated separately within each experimental background by subtracting the mean fluorescence of the corresponding untagged control. Data plotted in the figures show the mean background-subtracted fluorescence with 95% confidence intervals. Confidence intervals for differences between reporter-positive and no-reporter groups were calculated by combining the standard errors of the two independent groups.

### Use of artificial intelligence tools

ChatGPT (OpenAI) and Claude (Anthropic) was used to assist with language editing and improving the clarity and readability of the manuscript, and to assist in generating code used for data visualization and plotting. All AI-generated code was reviewed and validated by the authors. The scientific analyses, interpretation of the data and conclusions were performed and verified by the authors, who take full responsibility for the final manuscript.

## Acknowledgements

We are thankful to Cihan Elci for technical assistance and William Mair and Jordan Ward for sharing strains prior to publication. We acknowledge support by the Microscopy Imaging Center at the University of Bern. Some strains were provided by the CGC, which is funded by NIH Office of Research Infrastructure Programs (P40 OD010440). This research was funded in part by the Swiss National Science Foundation (SNSF) [grants PCEFP3_181204, 310030_219822, 320030L-227534]. For the purpose of open access, a CC BY public copyright licence is applied to any Author Accepted Manuscript (AAM) version arising from this submission.

## Data and resource availability

Code use for image analysis can be accessed at https://github.com/spsalmon/towbintools_pipeline.

## Author Contributions

**B.D.T** and **I.G.** formulated the research question, **I.G.** performed the experiments, analysed data, established image analysis workflows and drafted the manuscript. **K.S.** conducted preliminary research on the genome-wide RNAi candidate hits. **A.G.** and **D.C.B.** conducted the experimental aspect of the validation and suppressor RNAi screens, respectively. **D.B.** created the *vhp-1::AID* allele. **S.P.** established the image analysis workflow for longitudinal live imaging. **B.D.T.** supervised the project, guided experiments and data analysis, and edited the manuscript. All authors reviewed the manuscript.

