## Supplementary Information for "The MAPK phosphatase VHP-1 buffers pharynx-to-body proportions against tissue-specific growth imbalance in *C. elegans*"

- Supplementary Figures 1-7

- Supplementary Tables S1-S8

Supplementary Figures

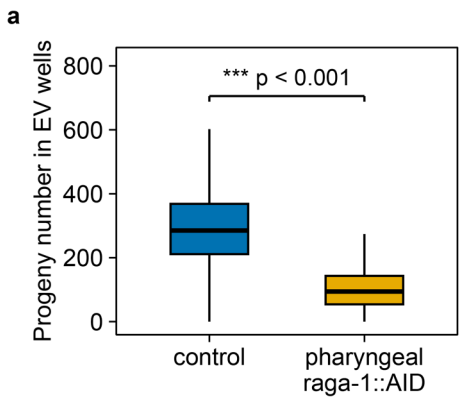

**Supplementary Figure 1. Worm count after 4 days is reduced upon pharyngeal RAGA-1 AID.** **a)** Worm count after four days per well in empty-vector (EV) RNAi conditions, in control (no RAGA-1 AID) and pharyngeal RAGA-1 AID animals. For both conditions $n = 1152$ wells.

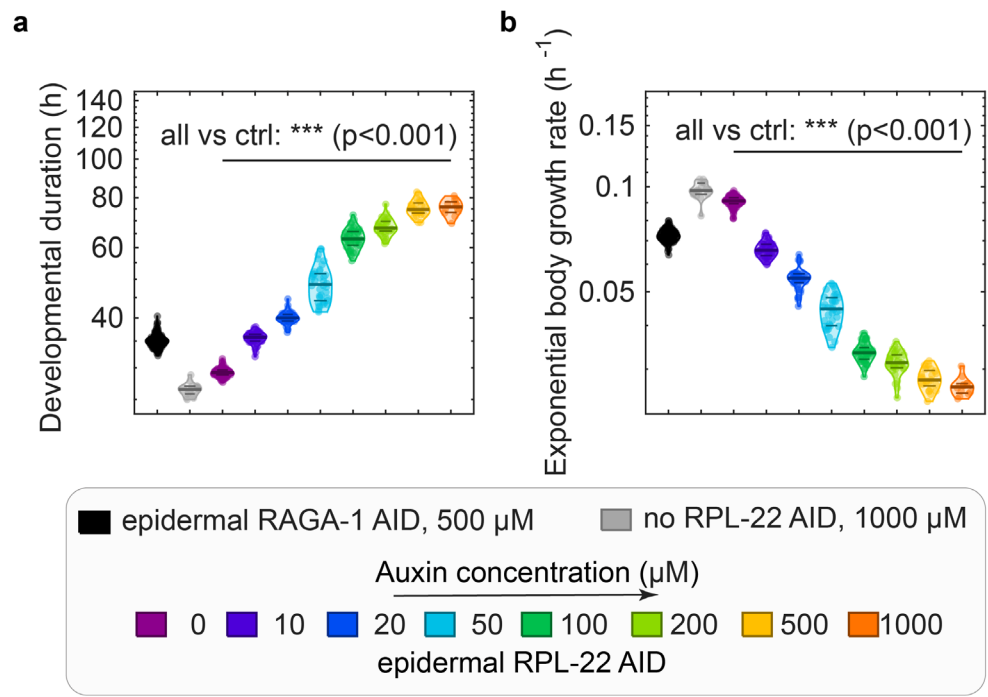

**Supplementary Figure 2. Titration of auxin concentration to reach equivalent growth rate for RPL-22 AID and RAGA-1 AID.** **a, b)** Developmental duration and exponential body growth rate from M1 to M4. Black indicates epidermal RAGA-1 AID animals treated with 500 $\mu\text{M}$ auxin, grey indicates the 1000 $\mu\text{M}$ control condition. Colored violins show epidermal RPL-22 AID animals exposed to increasing auxin concentrations (0–1000 $\mu\text{M}$). Individual animals are shown as points; horizontal lines indicate the 25th percentile, median, and 75th percentile. The RPL-22 AID titration conditions differed significantly from the control in both panels (\*\*\*, $p < 0.001$). Numbers of individual animals per condition ($n$) were 116, 18, 113, 53, 40, 38, 50, 35, 25, and 12, respectively.

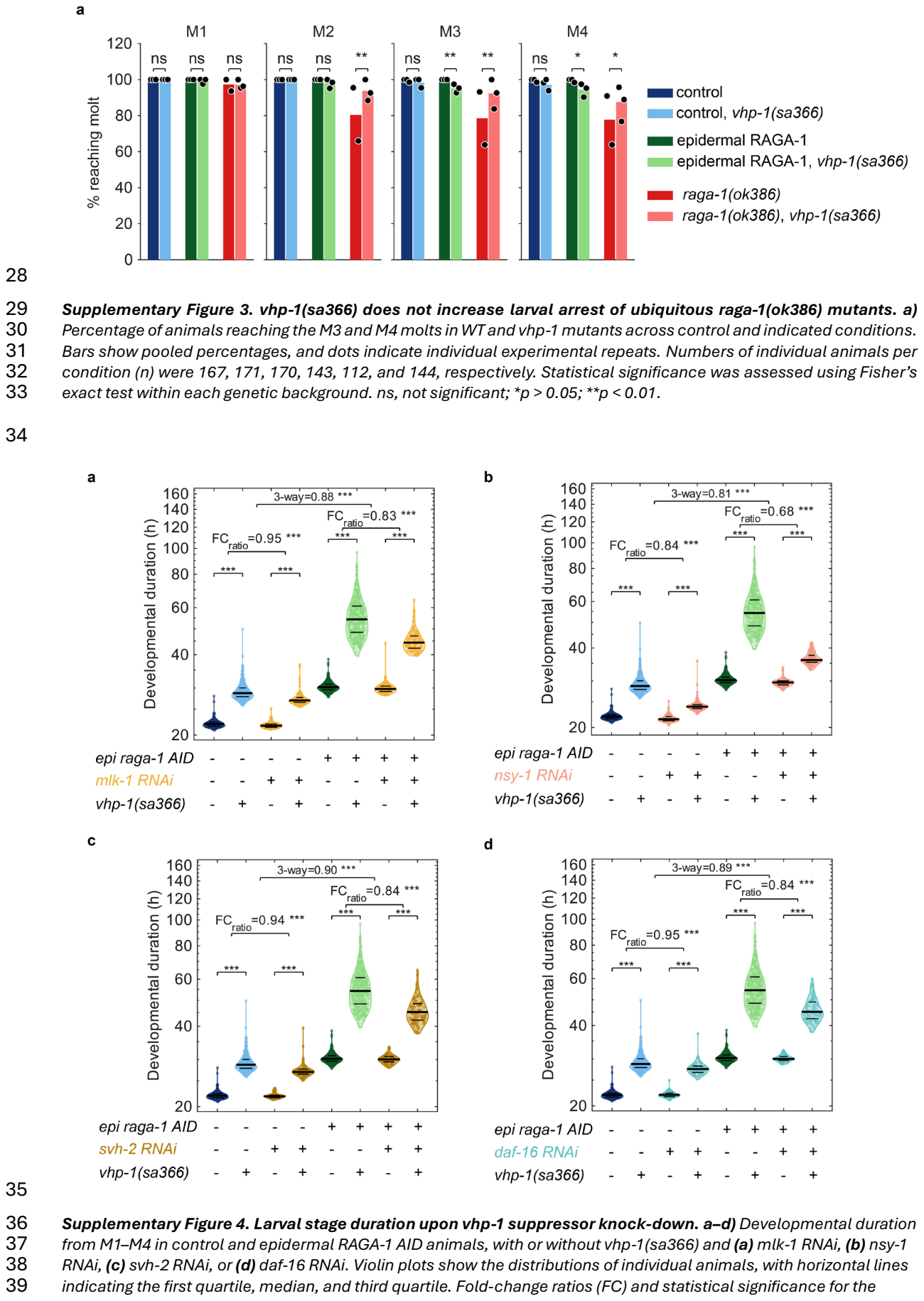

Supplementary Figure 3. vhp-1(sa366) does not increase larval arrest of ubiquitous raga-1(ok386) mutants. a) Percentage of animals reaching the M3 and M4 molts in WT and vhp-1 mutants across control and indicated conditions. Bars show pooled percentages, and dots indicate individual experimental repeats. Numbers of individual animals per condition (n) were 167, 171, 170, 143, 112, and 144, respectively. Statistical significance was assessed using Fisher’s exact test within each genetic background. ns, not significant; \*p > 0.05; \*\*p < 0.01.

Supplementary Figure 4. Larval stage duration upon vhp-1 suppressor knock-down. a–d) Developmental duration from M1–M4 in control and epidermal RAGA-1 AID animals, with or without vhp-1(sa366) and (a) mlk-1 RNAi, (b) nsy-1 RNAi, (c) svh-2 RNAi, or (d) daf-16 RNAi. Violin plots show the distributions of individual animals, with horizontal lines indicating the first quartile, median, and third quartile. Fold-change ratios (FC) and statistical significance for the indicated contrasts are shown above the plots. “3-way” denotes the interaction among epidermal RAGA-1 depletion, *vhp-1(sa366)*, and the indicated RNAi treatment. ns, not significant; \**p* < 0.05; \*\**p* < 0.01; \*\*\**p* < 0.001.

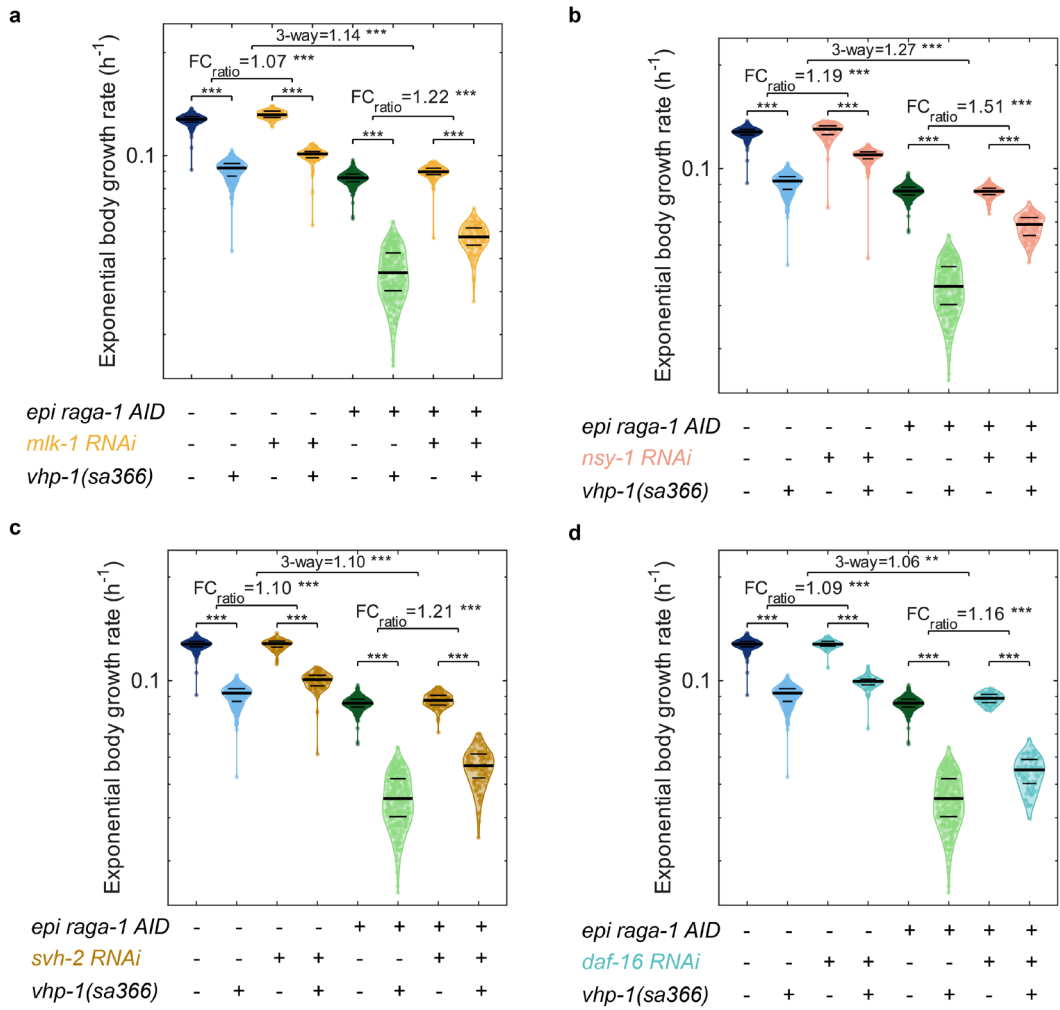

**Supplementary Figure 5. Exponential growth rate upon *vhp-1* suppressor knock-down.** a–d) Exponential growth rate from M1–M4 in control and epidermal RAGA-1 AID animals, with or without *vhp-1(sa366)* and (a) *mlk-1* RNAi, (b) *nsy-1* RNAi, (c) *svh-2* RNAi, or (d) *daf-16* RNAi. Violin plots show the distributions of individual animals, with horizontal lines indicating the first quartile, median, and third quartile. Fold-change ratios (FC) and statistical significance for the indicated contrasts are shown above the plots. “3-way” denotes the interaction among epidermal RAGA-1 depletion, *vhp-1(sa366)*, and the indicated RNAi treatment. ns, not significant; \**p* < 0.05; \*\**p* < 0.01; \*\*\**p* < 0.001.

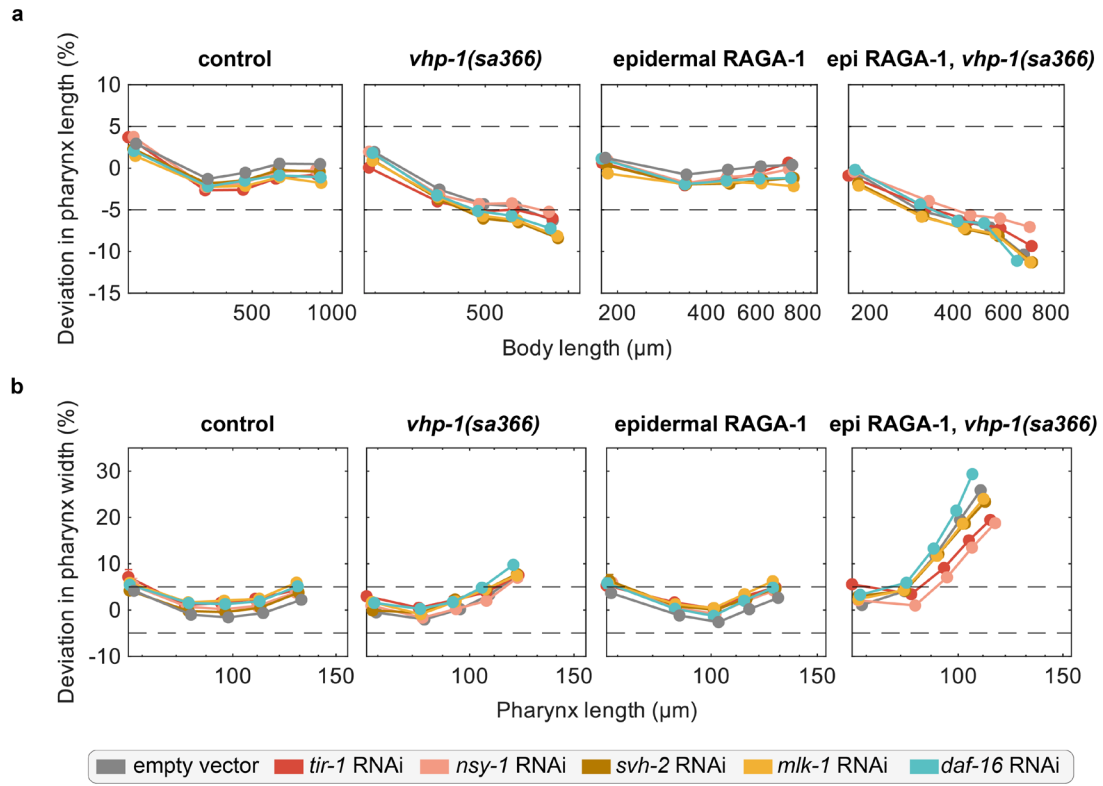

**Supplementary Figure 6. Pharynx width and length in *vhp-1* suppressors. a,b)** Mean percentage deviation in pharynx length (a) and pharynx width (b) in control, *vhp-1(sa366)*, epidermal RAGA-1 AID, and epidermal RAGA-1 AID; *vhp-1(sa366)* animals treated with empty vector or RNAi against *tir-1*, *nsy-1*, *svh-2*, *mlk-1*, or *daf-16*. Pharynx length deviation is plotted against body length, whereas pharynx width deviation is plotted against pharynx length. Each point represents the mean deviation at one ecdysis, and error bars indicate SEM across individuals. Separate control log-log models were fitted for each experimental set. Dashed horizontal lines indicate $\pm 5\%$ deviation from the control prediction.

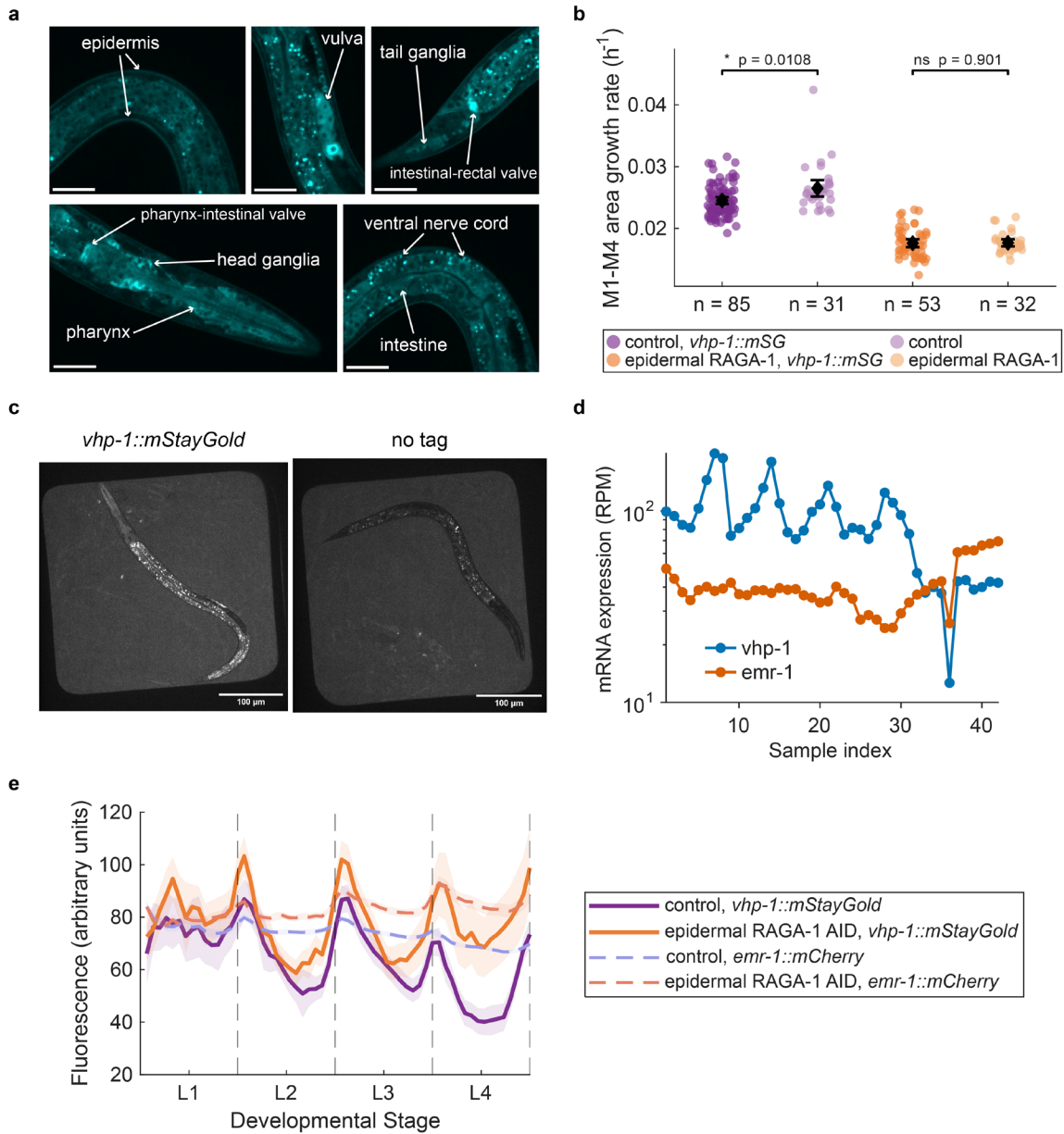

**Supplementary Figure 7. Characterization of VHP-1::mStayGold expression and reporter controls. a)** Enlarged images from the same animal expressing VHP-1::mStayGold as shown in Figure 6a. Expression is seen broadly across the animal, including in the epidermis, vulva, tail ganglia, intestinal-rectal valve, pharyngeal-intestinal valve, head ganglia, pharynx, ventral nerve cord, and intestine. Scale bar: 20 $\mu\text{m}$. **b)** Exponential body-area growth rate between M1-M4 in control and epidermal RAGA-1 animals with or without VHP-1::mStayGold tag. For each animal, body area was quantified as the median area across z-slices at each time point. Points represent individual animals; black diamonds indicate the mean, and error bars show 95% confidence intervals. Comparisons were performed using Welch's two-sample t-test. Growth rate is shown as area growth rate compared to volume growth rate in the main figure due to unreliable volume computation from transmitted light segmentation images used for this strain. **c)** Representative fluorescence images of *vhp-1::mStayGold* animals and matched untagged controls imaged in microchambers. Untagged animals were used to quantify background fluorescence arising from worm and bacterial autofluorescence, enabling background correction of the *vhp-1::mStayGold* signal in subsequent measurements. Scale bars, 100 $\mu\text{m}$. **d)** Time-course mRNA expression of *vhp-1* and *emr-1* from previously published RNA-sequencing data from the study of Meeuse et al., 2023. Expression values were normalized to sequencing depth and reported as reads per million (RPM); samples are shown in chronological order, and expression is plotted on a logarithmic scale. **e)** Fluorescence was quantified across larval development in control (N=4, n=108) and epidermal RAGA-1 AID (N=4, n=96) animals and aligned to individual developmental stages. For each animal and time point, fluorescence was averaged across the five brightest z-slices selected based on *vhp-1::mStayGold* intensity. *vhp-1::mStayGold* fluorescence (solid lines) was background-corrected by subtracting the mean signal of the corresponding no-reporter condition. *emr-1* *emr-1::mCherry* fluorescence (dashed lines) was corrected by subtracting a fixed camera offset of 100 fluorescence units.

*No autofluorescence background subtraction could be applied, as both strains expressed the EMR-1:mCherry tag.* *Lines show mean fluorescence and shaded regions indicate 95% confidence intervals. Dashed vertical lines indicate* *molt boundaries.*

#### Supplementary Tables

**Supplementary Table S1. Primary hits from the genome-wide RNAi screen.** FC, fold change in progeny output (observed pharyngeal RAGA-1 AID / value predicted from the matched control); P values are Benjamini-Hochberg FDR-adjusted one-sided binomial-test P values. Rows are ordered by increasing FC.

| RNAi target | WormBase ID | FC | Adjusted p-val |
| --- | --- | --- | --- |
| cank-26 | WBGene00019641 | 0.0097 | $2.01 \times 10^{-4}$ |
| str-22 pseudogene | WBGene00006089 | 0.01 | $2.01 \times 10^{-4}$ |
| btb-6 | WBGene00018438 | 0.011 | $2.01 \times 10^{-4}$ |
| srh-90 pseudogene | WBGene00005311 | 0.012 | 0.00239 |
| fbxb-98 | WBGene00015830 | 0.013 | $2.01 \times 10^{-4}$ |
| C08F1.6 | WBGene00015610 | 0.013 | $2.01 \times 10^{-4}$ |
| math-23 | WBGene00016718 | 0.014 | $2.01 \times 10^{-4}$ |
| bcmo-2 | WBGene00018755 | 0.015 | $2.01 \times 10^{-4}$ |
| F54D10.5 | WBGene00018806 | 0.015 | $2.01 \times 10^{-4}$ |
| fbxb-97 | WBGene00016885 | 0.016 | $2.01 \times 10^{-4}$ |
| F53C3.11 | WBGene00018754 | 0.016 | $2.01 \times 10^{-4}$ |
| C17A2.3 | WBGene00015871 | 0.016 | 0.00239 |
| C17A2.6 pseudogene | WBGene00015874 | 0.019 | 0.00239 |
| sea-2 | WBGene00004751 | 0.025 | 0.00239 |
| C52E2.4 or C52E2.5 | WBGene00016883 or<br>WBGene00016884 | 0.029 | $2.01 \times 10^{-4}$ |
| F52C6.2 | WBGene00018659 | 0.032 | $2.01 \times 10^{-4}$ |
| his-61 | WBGene00001935 | 0.034 | $2.01 \times 10^{-4}$ |
| Y50E8A.11 | WBGene00013054 | 0.037 | $2.01 \times 10^{-4}$ |
| acs-6 | WBGene00022849 | 0.041 | 0.00239 |
| col-119 | WBGene00000693 | 0.042 | $2.01 \times 10^{-4}$ |
| math-4 | WBGene00015609 | 0.05 | $2.01 \times 10^{-4}$ |
| snfc-5 | WBGene00011111 | 0.055 | $2.01 \times 10^{-4}$ |
| gob-1 | WBGene00001649 | 0.069 | $2.01 \times 10^{-4}$ |
| cdc-14 | WBGene00000383 | 0.07 | $2.01 \times 10^{-4}$ |
| F42F12.4 | WBGene00009636 | 0.075 | $2.01 \times 10^{-4}$ |
| Y47D3B.3 | WBGene00012940 | 0.08 | $2.01 \times 10^{-4}$ |
| zyg-11 | WBGene00006996 | 0.086 | $2.01 \times 10^{-4}$ |
| mex-5 | WBGene00003230 | 0.09 | $2.01 \times 10^{-4}$ |
| syx-4 | WBGene00006374 | 0.091 | $2.01 \times 10^{-4}$ |
| bus-8B | WBGene00044623 | 0.092 | $2.01 \times 10^{-4}$ |
| C42C1.3 | WBGene00016581 | 0.093 | $2.01 \times 10^{-4}$ |
| dma-1 | WBGene00011345 | 0.097 | $2.01 \times 10^{-4}$ |
| mei-2 | WBGene00003184 | 0.1 | $2.01 \times 10^{-4}$ |
| math-5 | WBGene00015834 | 0.104 | $2.01 \times 10^{-4}$ |
| glp-1 | WBGene00001609 | 0.108 | 0.00239 |
| cap-1 | WBGene00000292 | 0.11 | $2.01 \times 10^{-4}$ |
| math-17 pseudogene | WBGene00015833 | 0.114 | $2.01 \times 10^{-4}$ |
| nhr-75 pseudogene | WBGene00003665 | 0.115 | 0.00239 |
| clec-2 | WBGene00015198 | 0.122 | $2.01 \times 10^{-4}$ |
| his-65 | WBGene00001939 | 0.123 | 0 |
| unc-15 | WBGene00006754 | 0.128 | $2.01 \times 10^{-4}$ |
| F55C12.19 | WBGene00271817 | 0.137 | $2.01 \times 10^{-4}$ |
| lit-1 | WBGene00003048 | 0.137 | $2.01 \times 10^{-4}$ |
| math-15 | WBGene00015829 | 0.137 | $2.01 \times 10^{-4}$ |
| ptc-2 pseudogene | WBGene00004209 | 0.144 | $2.01 \times 10^{-4}$ |
| C17A2.7 | WBGene00015875 | 0.147 | $2.01 \times 10^{-4}$ |
| hgrs-1 | WBGene00004101 | 0.147 | $1.03 \times 10^{-6}$ |
| lrr-1 | WBGene00018016 | 0.152 | $2.01 \times 10^{-4}$ |
| Y46H3C.e | — | 0.155 | $2.01 \times 10^{-4}$ |
| edg-1 | WBGene00007109 | 0.155 | $2.01 \times 10^{-4}$ |
| nhr-119 | WBGene00003709 | 0.155 | $2.01 \times 10^{-4}$ |
| sru-2 | WBGene00005665 | 0.157 | $2.01 \times 10^{-4}$ |
| W06A11.1 | WBGene00021052 | 0.161 | 0.00239 |
| gfi-1 | WBGene00001581 | 0.163 | $4.27 \times 10^{-12}$ |
| pole-2 | WBGene00017237 | 0.164 | $2.01 \times 10^{-4}$ |
| skr-2 | WBGene00004808 | 0.165 | $2.01 \times 10^{-4}$ |
| oac-45 | WBGene00011656 | 0.165 | $2.01 \times 10^{-4}$ |
| his-46 | WBGene00001920 | 0.174 | 0 |
| egg-6 | WBGene00010621 | 0.181 | $2.01 \times 10^{-4}$ |
| B0280.9 | WBGene00015104 | 0.182 | $2.01 \times 10^{-4}$ |
| ccdc-55 | WBGene00007627 | 0.187 | $2.01 \times 10^{-4}$ |
| srw-79 | WBGene00005826 | 0.189 | $2.01 \times 10^{-4}$ |
| bus-24 | WBGene00012433 | 0.19 | $2.01 \times 10^{-4}$ |
| clec-123 | WBGene00021288 | 0.191 | $2.01 \times 10^{-4}$ |
| hlh-11 | WBGene00001955 | 0.191 | $2.01 \times 10^{-4}$ |
| ZC239.14 | WBGene00022572 | 0.193 | $2.01 \times 10^{-4}$ |
| pro-1 | WBGene00004185 | 0.194 | $2.01 \times 10^{-4}$ |
| rpa-1 | WBGene00017546 | 0.195 | $2.01 \times 10^{-4}$ |
| C16C8.5 | WBGene00015843 | 0.196 | 0.00239 |
| R03H10.4 | WBGene00019856 | 0.196 | 0.00239 |
| sna-2 | WBGene00011747 | 0.197 | $2.01 \times 10^{-4}$ |
| cpar-1 | WBGene00010036 | 0.199 | $2.01 \times 10^{-4}$ |
| vhp-1 | WBGene00006923 | 0.199 | 0.00239 |
| C08F1.8 | WBGene00015611 | 0.199 | $2.01 \times 10^{-4}$ |
| T10D4.1 | WBGene00020406 | 0.2 | $2.01 \times 10^{-4}$ |

**Supplementary Table S2. Validation screen hits.** Values are $\log_2$ fold changes and Benjamini-Hochberg-adjusted one-sided binomial-test *P* values. For tissue-specific hits, values are shown only for the background in which the clone met the validation criteria.

| Gene | Pharyngeal $\log_2$FC | Pharyngeal p-val | Epidermal $\log_2$FC | Epidermal p-val |
| --- | --- | --- | --- | --- |
| **Shared hits** |  |  |  |  |
| *cap-1* | -2.23 | $1.77 \times 10^{-9}$ | -1.22 | $3.08 \times 10^{-5}$ |
| *cks-1* | -1.70 | $7.81 \times 10^{-4}$ | -1.28 | $6.56 \times 10^{-11}$ |
| *cpar-1* | -1.52 | $3.53 \times 10^{-6}$ | -1.74 | $6.56 \times 10^{-11}$ |
| *gob-1* | -1.43 | $4.51 \times 10^{-12}$ | -1.01 | $5.04 \times 10^{-8}$ |
| *pole-2* | -1.12 | $5.03 \times 10^{-8}$ | -1.30 | $3.08 \times 10^{-5}$ |
| *skr-2* | -1.52 | $1.77 \times 10^{-9}$ | -1.52 | $5.04 \times 10^{-8}$ |
| ***vhp-1*** | -1.39 | 0.00407 | -1.72 | $1.65 \times 10^{-6}$ |
| Y11D7A.9 | -1.85 | $3.82 \times 10^{-11}$ | -1.53 | $5.04 \times 10^{-8}$ |
| **Pharyngeal-specific hits** |  |  |  |  |
| *col-119* | -1.13 | 0.00184 | — | — |
| *edg-1* | -1.36 | $1.13 \times 10^{-4}$ | — | — |
| *mei-2* | -1.15 | $3.53 \times 10^{-6}$ | — | — |
| *nhr-261* | -1.02 | $2.78 \times 10^{-10}$ | — | — |
| *unc-15* | -1.90 | $1.22 \times 10^{-5}$ | — | — |
| Y14H12B.1 | -1.13 | $2.78 \times 10^{-10}$ | — | — |
| **Epidermal-specific hits** |  |  |  |  |
| *bub-3* | — | — | -1.04 | $1.65 \times 10^{-6}$ |
| C42C1.3 | — | — | -2.28 | $7.57 \times 10^{-10}$ |
| F55C12.3 | — | — | -1.42 | $7.03 \times 10^{-8}$ |
| *lrr-1* | — | — | -2.95 | $3.91 \times 10^{-14}$ |
| *mex-5* | — | — | -1.33 | $6.56 \times 10^{-11}$ |
| *pro-1* | — | — | -1.96 | $4.18 \times 10^{-12}$ |
| *pygl-1* | — | — | -1.03 | $7.61 \times 10^{-6}$ |
| Y19D2B.1 | — | — | -1.44 | $4.18 \times 10^{-12}$ |

**Supplementary Table S3. Sample sizes and pairwise statistics for Figure 2.** Values are EV/vhp-1(RNAi). N, independent day-to-day repeats; n, animals included. For b/e, the result is animals reaching the indicated molt (EV vs RNAi; two-sided Fisher exact test). For c/d, FC = median RNAi/median EV (two-sided Wilcoxon rank-sum test); c, developmental duration; d, exponential body growth rate.

| Panel | Background | Stage | N; n<br>(EV/RNAi) | Result | P |
| --- | --- | --- | --- | --- | --- |
| **b** | Control | M3 | 4/4; 180/180 | 180/180 vs 176/180 | 0.123 |
| | | M4 | 4/4; 180/180 | 180/180 vs 85/180 | $1.73 \times 10^{-36}$ |
| | Epidermal<br>RAGA-1 | M3 | 4/4; 208/194 | 208/208 vs 104/194 | $2.63 \times 10^{-35}$ |
| | | M4 | 4/4; 208/194 | 208/208 vs 17/194 | $4.56 \times 10^{-95}$ |
|  | Pharyngeal<br>RAGA-1 | M3 | 2/2; 86/90 | 86/86 vs 89/90 | 1 |
| | | M4 | 2/2; 86/90 | 85/86 vs 9/90 | $1.64 \times 10^{-38}$ |
| **c** | Control | M1-M3 | 4/4; 181/171 | FC 1.368 | $6.64 \times 10^{-59}$ |
| | Epidermal<br>RAGA-1 | M1-M3 | 4/4; 208/104 | FC 1.602 | $5.45 \times 10^{-47}$ |
| | Pharyngeal<br>RAGA-1 | M1-M3 | 2/2; 86/89 | FC 1.380 | $9.82 \times 10^{-30}$ |
| **d** | Control | M1-M3 | 4/4; 181/176 | FC 0.660 | $5.29 \times 10^{-60}$ |
| | Epidermal<br>RAGA-1 | M1-M3 | 4/4; 208/104 | FC 0.494 | $5.29 \times 10^{-47}$ |
| | Pharyngeal<br>RAGA-1 | M1-M3 | 2/2; 84/87 | FC 0.675 | $4.04 \times 10^{-28}$ |
| **e** | Control | M1 | 2/2; 114/98 | 113/114 vs 98/98 | 1 |
|  |  | M2 | 2/2; 114/98 | 111/114 vs 98/98 | 0.251 |
|  |  | M3 | 2/2; 114/98 | 108/114 vs 97/98 | 0.126 |
| | | M4 | 2/2; 114/98 | 108/114 vs 60/98 | $1.28 \times 10^{-9}$ |
| | Epidermal<br>RPL-22 | M1 | 2/2; 76/90 | 76/76 vs 76/90 | $1.23 \times 10^{-4}$ |
| | | M2 | 2/2; 76/90 | 75/76 vs 5/90 | $6.44 \times 10^{-40}$ |
| | | M3 | 2/2; 76/90 | 75/76 vs 0/90 | $2.83 \times 10^{-47}$ |
| | | M4 | 2/2; 76/90 | 75/76 vs 0/90 | $2.83 \times 10^{-47}$ |

**Supplementary Table S4. Sample sizes and pairwise statistics for Figure 3b.** Figure 3b uses the same independent experiments as Figure 2b–d. Each stage cell shows *n* (mock RNAi/vhp-1 RNAi) on the first line and the *P* value on the second line. *P* values are from two-sided Wilcoxon rank-sum tests comparing per-animal deviations between mock RNAi and vhp-1(RNAi) at the indicated ecdysis.

| Group | Trait | Hatch | M1 | M2 | M3 | M4 |
| --- | --- | --- | --- | --- | --- | --- |
| Control | Volume | 109/128<br>0.696 | 182/180<br>9.60E-15 | 182/178<br>4.69E-06 | 182/177<br>0.00456 | 180/85<br>2.51E-16 |
|  | Width | 109/128<br>9.75E-08 | 182/180<br>2.57E-04 | 182/178<br>0.864 | 182/177<br>2.72E-06 | 180/85<br>0.0011 |
|  | Length | 109/128<br>0.00285 | 182/180<br>5.85E-13 | 182/178<br>2.21E-13 | 182/177<br>2.55E-09 | 180/85<br>5.25E-11 |
| Epidermal RAGA-1 AID | Volume | 123/125<br>0.0115 | 208/192<br>1.31E-24 | 208/190<br>9.68E-35 | 208/104<br>9.80E-41 | 208/17<br>1.03E-11 |
|  | Width | 123/125<br>3.45E-12 | 208/192<br>1.44E-05 | 208/190<br>5.07E-19 | 208/104<br>4.49E-29 | 208/17<br>8.57E-06 |
|  | Length | 123/125<br>0.31 | 208/192<br>1.51E-32 | 208/190<br>3.43E-47 | 208/104<br>1.41E-46 | 208/17<br>7.43E-12 |
| Pharyngeal RAGA-1 AID | Volume | 38/38<br>0.15 | 86/89<br>0.304 | 86/89<br>0.389 | 86/89<br>0.00482 | 85/9<br>0.136 |
|  | Width | 38/38<br>4.16E-05 | 86/89<br>0.00327 | 86/89<br>0.55 | 86/89<br>0.0206 | 85/9<br>0.0299 |
|  | Length | 38/38<br>0.049 | 86/89<br>0.0313 | 86/89<br>0.233 | 86/89<br>0.00181 | 85/9<br>0.00232 |

**Cell format: n mock/RNAi; P.** Volume, width, and length refer to pharyngeal traits. *n* is the number of animals with valid measurements for that trait and ecdysis.

**Supplementary Table S5. Sample sizes and pairwise statistics for Figure 4 and Supplemental Figure 3.** All comparisons are wild-type *vhp-1* versus *vhp-1(sa366)* within the indicated *RAGA-1* background; *N* = 3 independent experiments per group. Exact *P* values are shown.

**Figure 4b-c**

| Group | Duration<br>n | FC | P | Growth<br>n | FC | P |
| --- | --- | --- | --- | --- | --- | --- |
| Control | 166/166 | 1.498 | 6.25E-56 | 165/166 | 0.581 | 9.19E-56 |
| Epidermal<br>RAGA-1 AID | 165/135 | 1.810 | 3.32E-50 | 166/135 | 0.501 | 2.46E-50 |
| *raga-1(ok386)* | 83/122 | 1.179 | 1.02E-23 | 83/124 | 0.729 | 6.03E-30 |

**Figure 4d**

| Group | Trait | Hatch | M1 | M2 | M3 | M4 |
| --- | --- | --- | --- | --- | --- | --- |
| Control | Volume | 119/92<br>0.0548 | 167/171<br>0.397 | 167/171<br>4.56E-46 | 166/168<br>8.42E-52 | 166/166<br>7.02E-51 |
|  | Width | 119/92<br>7.99E-08 | 167/171<br>6.65E-04 | 167/171<br>1.46E-43 | 166/168<br>1.43E-54 | 166/166<br>1.81E-54 |
|  | Length | 119/92<br>0.255 | 167/171<br>3.02E-11 | 167/171<br>8.17E-09 | 166/168<br>3.84E-39 | 166/166<br>4.03E-46 |
| Epidermal<br>RAGA-1 AID | Volume | 119/97<br>0.0226 | 182/180<br>0.00698 | 182/177<br>1.60E-56 | 181/167<br>1.51E-50 | 169/135<br>4.11E-46 |
|  | Width | 119/97<br>1.62E-06 | 182/180<br>2.79E-41 | 182/177<br>3.60E-60 | 181/167<br>1.98E-58 | 169/135<br>9.97E-51 |
|  | Length | 119/97<br>0.00453 | 182/180<br>3.84E-53 | 182/177<br>9.13E-40 | 181/167<br>1.86E-50 | 169/135<br>3.08E-49 |
| *raga-1(ok386)* | Volume | 75/102<br>0.664 | 110/140<br>4.21E-14 | 91/135<br>2.13E-21 | 89/133<br>5.18E-24 | 87/126<br>7.16E-21 |
|  | Width | 75/102<br>0.00187 | 110/140<br>0.0306 | 91/135<br>1.60E-12 | 89/133<br>2.00E-21 | 87/126<br>1.11E-22 |
|  | Length | 75/102<br>0.0082 | 110/140<br>0.0883 | 91/135<br>0.147 | 89/133<br>5.55E-07 | 87/126<br>2.36E-16 |

**Supplementary Figure 3**

| Background | Total n | M1 | M2 | M3 | M4 |
| --- | --- | --- | --- | --- | --- |
| Control | 167/171 | 167/171<br>1 | 167/171<br>1 | 166/168<br>0.623 | 166/166<br>0.215 |
| Epidermal<br>RAGA-1 AID | 170/143 | 170/142<br>0.457 | 170/140<br>0.0943 | 170/136<br>0.00383 | 169/135<br>0.013 |
| *raga-1(ok386)* | 112/144 | 109/140<br>1 | 90/135<br>0.00167 | 88/133<br>0.00174 | 87/126<br>0.0435 |

**b-c:** *n* = animals after endpoint-specific exclusions and 3×IQR filtering on log-transformed values; FC = median(*sa366*)/median(WT); *P*, two-sided Wilcoxon rank-sum. **d:** stage cells show *n* WT/*sa366* and *P* from two-sided Wilcoxon rank-sum tests of per-animal deviations; no 3×IQR filter. **Supplemental Figure 3:** Total *n* = counted WT/*sa366*; stage cells show number reaching the molt WT/*sa366* and Fisher exact *P*.

**Supplementary Table S6. RNAi suppressor-screen hits from Figure 5b.** FC is mean progeny per adult relative to EV. Hits met adjusted $P < 5 \times 10^{-6}$ and $FC > 4$. The screen pooled two independent repeats; all listed hits had $n = 12$ observations. RNAi clone column states the Ahringer or Vidal library position of each clone.

| Gene | RNAi clone | FC | Adjusted P |
| --- | --- | --- | --- |
| *tir-1* | *tir-1* 72/G10 | 8.948 | 2.73E-7 |
| *svh-2* | *svh-2* 191/A12 | 8.380 | 2.73E-7 |
| *nsy-1* | *nsy-1* 45/G01 | 7.656 | 1.20E-6 |
| *daf-16* | *daf-16* 10020@B8 | 6.425 | 7.47E-7 |
| *mlk-1* | *mlk-1* 141/C01 | 6.136 | 3.18E-7 |
| *jun-1* | *jun-1* 10113@F2 | 5.902 | 7.28E-7 |
| *zak-1* | *zak-1* 79/E02 | 5.115 | 2.61E-6 |
| *mek-1* | *mek-1* 192/F02 | 5.082 | 1.09E-6 |
| *pmk-2* | *pmk-2* 11049@F11 | 4.817 | 1.64E-6 |
| *wnk-1* | *wnk-1* 11024@F04 | 4.204 | 2.60E-6 |
| *skn-1* | *skn-1* 10144@G11 | 4.173 | 3.77E-6 |

**Supplementary Table S7. Sample sizes and statistical results for Figure 5d–h and Supplemental Figures 4–6.** Unless noted otherwise, $P$ values are from two-sided Wilcoxon rank-sum tests.

###### Longitudinal sample sizes

| RNAi | No RAGA-1<br>WT | No RAGA-1<br>*sa366* | Epi RAGA-1 AID<br>WT | Epi RAGA-1 AID<br>*sa366* |
| --- | --- | --- | --- | --- |
| EV | 10/384 | 10/348 | 10/381 | 10/392 |
| *tir-1* | 3/114 | 3/122 | 3/121 | 3/144 |
| *nsy-1* | 3/101 | 3/117 | 3/118 | 3/149 |
| *svh-2* | 3/92 | 3/115 | 3/120 | 3/121 |
| *mlk-1* | 3/99 | 3/111 | 3/126 | 3/136 |
| *daf-16* | 2/77 | 2/73 | 2/86 | 2/98 |

Cells show N/n, where N is the number of independent experiments and n is the number of animals contributing to the M1–M4 duration/growth analyses.

###### Figure 5f–g: direct suppressor comparison in epidermal RAGA-1 AID; *vhp-1(sa366)*

| RNAi | Duration FC | P | Growth FC | P |
| --- | --- | --- | --- | --- |
| *tir-1* | 0.684 | 6.28E-68 | 1.483 | 3.63E-67 |
| *svh-2* | 0.836 | 5.23E-29 | 1.244 | 2.05E-31 |
| *nsy-1* | 0.666 | 3.37E-72 | 1.512 | 4.43E-71 |
| *mlk-1* | 0.821 | 1.37E-39 | 1.271 | 5.60E-44 |
| *daf-16* | 0.832 | 1.51E-25 | 1.210 | 5.12E-21 |

FC = median(RNAi)/median(EV).

###### Figure 5d–e and Supplemental Figures 4–5: WT versus *vhp-1(sa366)* within each condition

| RNAi | Endpoint | EV<br>no AID | RNAi<br>no AID | EV<br>AID | RNAi<br>AID |
| --- | --- | --- | --- | --- | --- |
| *tir-1* | Duration | 1.303<br>1.49E-120 | 1.140<br>2.01E-37 | 1.791<br>6.57E-128 | 1.276<br>1.30E-44 |
| *tir-1* | Growth rate | 0.719<br>3.01E-120 | 0.846<br>2.00E-38 | 0.529<br>6.59E-128 | 0.748<br>1.22E-44 |
| *nsy-1* | Duration | 1.303<br>1.49E-120 | 1.116<br>5.43E-35 | 1.791<br>6.57E-128 | 1.216<br>1.28E-44 |
| *nsy-1* | Growth rate | 0.719<br>3.01E-120 | 0.843<br>1.50E-34 | 0.529<br>6.59E-128 | 0.801<br>2.25E-44 |
| *svh-2* | Duration | 1.303<br>1.49E-120 | 1.237<br>4.74E-35 | 1.791<br>6.57E-128 | 1.502<br>4.90E-41 |
| *svh-2* | Growth rate | 0.719<br>3.01E-120 | 0.787<br>4.76E-35 | 0.529<br>6.59E-128 | 0.645<br>4.91E-41 |
| *mlk-1* | Duration | 1.303<br>1.49E-120 | 1.246<br>7.55E-36 | 1.791<br>6.57E-128 | 1.490<br>8.68E-44 |
| *mlk-1* | Growth rate | 0.719<br>3.01E-120 | 0.770<br>7.60E-36 | 0.529<br>6.59E-128 | 0.644<br>1.12E-43 |
| *daf-16* | Duration | 1.303<br>1.49E-120 | 1.250<br>4.23E-26 | 1.791<br>6.57E-128 | 1.500<br>1.45E-31 |
| *daf-16* | Growth rate | 0.719<br>3.01E-120 | 0.779<br>4.47E-26 | 0.529<br>6.59E-128 | 0.620<br>1.45E-31 |

Each condition cell shows FC and exact P. FC = median(sa366)/median(WT).

###### Interaction contrasts

| RNAi | Endpoint | FC ratio<br>no AID | FC ratio<br>AID | 3-way ratio |
| --- | --- | --- | --- | --- |
| *tir-1* | Duration | 0.857<br>7.44E-35 | 0.713<br>5.41E-154 | 0.832<br>9.02E-27 |
| *tir-1* | Growth rate | 1.189<br>1.01E-37 | 1.411<br>2.92E-140 | 1.187<br>1.63E-20 |
| *nsy-1* | Duration | 0.844<br>1.70E-39 | 0.682<br>1.42E-190 | 0.807<br>1.74E-34 |
| *nsy-1* | Growth rate | 1.187<br>4.79E-33 | 1.510<br>2.43E-179 | 1.272<br>6.12E-35 |
| *svh-2* | Duration | 0.937<br>1.66E-6 | 0.841<br>3.37E-41 | 0.897<br>5.08E-9 |
| *svh-2* | Growth rate | 1.103<br>4.78E-11 | 1.214<br>9.64E-43 | 1.101<br>2.31E-6 |
| *mlk-1* | Duration | 0.947<br>3.97E-5 | 0.830<br>2.18E-51 | 0.876<br>1.69E-13 |
| *mlk-1* | Growth rate | 1.069<br>2.70E-6 | 1.219<br>1.39E-49 | 1.140<br>1.52E-11 |
| *daf-16* | Duration | 0.947<br>3.92E-4 | 0.841<br>1.35E-33 | 0.888<br>1.33E-8 |
| *daf-16* | Growth rate | 1.088<br>3.76E-7 | 1.157<br>3.19E-21 | 1.063<br>0.00682 |

Cells show ratio and exact model-contrast P. The 3-way term tests whether the *vhp-1* × RNAi FC ratio differs with epidermal RAGA-1 depletion.

**Figure 5h / Supplemental Figure 6: Control**

EV reference n (Hatch–M4): 126, 393, 393, 393, 384. Stage cells show n for the RNAi condition and exact P versus the matched EV group.

| RNAi | Trait | Hatch | M1 | M2 | M3 | M4 |
| --- | --- | --- | --- | --- | --- | --- |
| *tir-1* | volume | 44<br>0.544 | 114<br>2.66E-8 | 114<br>3.93E-4 | 114<br>0.176 | 114<br>3.08E-7 |
| *tir-1* | width | 44<br>0.128 | 114<br>9.34E-8 | 114<br>1.26E-17 | 114<br>3.90E-19 | 114<br>5.42E-8 |
| *tir-1* | length | 44<br>0.253 | 114<br>2.14E-11 | 114<br>1.50E-22 | 114<br>8.72E-22 | 114<br>3.73E-11 |
| *nsy-1* | volume | 30<br>0.622 | 101<br>4.38E-7 | 101<br>4.98E-16 | 101<br>1.72E-11 | 101<br>5.17E-8 |
| *nsy-1* | width | 30<br>0.697 | 101<br>1.04E-5 | 101<br>2.52E-6 | 101<br>8.49E-4 | 101<br>0.00428 |
| *nsy-1* | length | 30<br>0.292 | 101<br>6.92E-7 | 101<br>1.97E-8 | 101<br>3.20E-7 | 101<br>2.68E-5 |
| *svh-2* | volume | 59<br>0.0319 | 93<br>0.0266 | 93<br>0.0228 | 93<br>0.938 | 92<br>0.558 |
| *svh-2* | width | 59<br>0.725 | 93<br>0.0602 | 93<br>0.00654 | 93<br>0.00112 | 92<br>4.17E-5 |
| *svh-2* | length | 59<br>0.215 | 93<br>0.00875 | 93<br>1.87E-5 | 93<br>2.47E-4 | 92<br>8.79E-5 |
| *mlk-1* | volume | 65<br>0.379 | 122<br>4.32E-5 | 122<br>1.71E-4 | 122<br>0.0258 | 99<br>7.57E-5 |
| *mlk-1* | width | 65<br>0.101 | 122<br>8.47E-14 | 122<br>2.61E-24 | 122<br>2.04E-20 | 99<br>2.13E-21 |
| *mlk-1* | length | 65<br>7.22E-4 | 122<br>2.96E-6 | 122<br>1.58E-15 | 122<br>7.52E-19 | 99<br>3.52E-25 |
| *daf-16* | volume | 47<br>0.172 | 92<br>5.05E-4 | 92<br>0.0924 | 92<br>0.0485 | 77<br>0.775 |
| *daf-16* | width | 47<br>0.0902 | 92<br>9.37E-11 | 92<br>6.99E-13 | 92<br>5.93E-12 | 77<br>6.97E-10 |
| *daf-16* | length | 47<br>0.146 | 92<br>1.14E-5 | 92<br>2.74E-5 | 92<br>3.23E-11 | 77<br>1.34E-13 |

**Figure 5h / Supplemental Figure 6: *vhp-1(sa366)***

EV reference n (Hatch–M4): 245, 356, 354, 353, 348. Stage cells show n for the RNAi condition and exact P versus the matched EV group.

| RNAi | Trait | Hatch | M1 | M2 | M3 | M4 |
| --- | --- | --- | --- | --- | --- | --- |
| *tir-1* | volume | 89<br>0.0259 | 130<br>3.35E-11 | 124<br>3.75E-6 | 123<br>3.90E-5 | 122<br>1.68E-10 |
| *tir-1* | width | 89<br>3.92E-7 | 130<br>1.25E-11 | 124<br>1.01E-6 | 123<br>0.135 | 122<br>0.979 |
| *tir-1* | length | 89<br>4.89E-6 | 130<br>1.34E-11 | 124<br>1.47E-4 | 123<br>0.143 | 122<br>0.144 |
| *nsy-1* | volume | 65<br>0.169 | 118<br>1.33E-13 | 118<br>9.32E-20 | 118<br>2.14E-24 | 117<br>8.94E-8 |
| *nsy-1* | width | 65<br>0.0481 | 118<br>0.217 | 118<br>0.574 | 118<br>0.0142 | 117<br>0.313 |
| *nsy-1* | length | 65<br>0.527 | 118<br>6.61E-4 | 118<br>0.754 | 118<br>0.124 | 117<br>2.71E-6 |
| *svh-2* | volume | 81<br>0.0789 | 115<br>6.86E-4 | 115<br>0.00205 | 115<br>5.24E-7 | 115<br>3.23E-22 |
| *svh-2* | width | 81<br>0.711 | 115<br>1.04E-4 | 115<br>2.63E-9 | 115<br>4.04E-4 | 115<br>0.217 |
| *svh-2* | length | 81<br>0.0215 | 115<br>6.11E-10 | 115<br>2.10E-14 | 115<br>7.33E-17 | 115<br>6.35E-20 |
| *mlk-1* | volume | 76<br>0.714 | 111<br>0.00141 | 111<br>0.00282 | 111<br>4.03E-7 | 111<br>3.75E-27 |
| *mlk-1* | width | 76<br>0.0262 | 111<br>7.24E-4 | 111<br>1.17E-7 | 111<br>1.72E-4 | 111<br>0.448 |
| *mlk-1* | length | 76<br>0.0562 | 111<br>1.22E-5 | 111<br>5.80E-10 | 111<br>1.92E-13 | 111<br>5.92E-18 |
| *daf-16* | volume | 66<br>0.0613 | 75<br>0.0551 | 75<br>1.80E-12 | 75<br>3.08E-20 | 73<br>1.76E-26 |
| *daf-16* | width | 66<br>0.00151 | 75<br>9.29E-8 | 75<br>1.24E-4 | 75<br>4.30E-5 | 73<br>3.25E-6 |
| *daf-16* | length | 66<br>0.948 | 75<br>0.00169 | 75<br>8.20E-4 | 75<br>1.22E-5 | 73<br>1.62E-4 |

### Figure 5h / Supplemental Figure 6: Epidermal RAGA-1 AID

EV reference n (Hatch–M4): 153, 402, 402, 402, 381. Stage cells show n for the RNAi condition and exact P versus the matched EV group.

| RNAi | Trait | Hatch | M1 | M2 | M3 | M4 |
| --- | --- | --- | --- | --- | --- | --- |
| *tir-1* | volume | 51<br>0.341 | 122<br>0.0931 | 122<br>0.0529 | 122<br>0.0764 | 121<br>0.0392 |
| *tir-1* | width | 51<br>0.0477 | 122<br>3.84E-19 | 122<br>4.62E-23 | 122<br>3.48E-20 | 121<br>2.54E-13 |
| *tir-1* | length | 51<br>0.317 | 122<br>1.01E-15 | 122<br>1.65E-10 | 122<br>1.96E-5 | 121<br>0.188 |
| *nsy-1* | volume | 42<br>0.395 | 133<br>3.60E-7 | 133<br>3.42E-7 | 133<br>0.00517 | 118<br>0.346 |
| *nsy-1* | width | 42<br>0.0257 | 133<br>5.38E-6 | 133<br>7.84E-9 | 133<br>1.03E-6 | 118<br>1.58E-4 |
| *nsy-1* | length | 42<br>0.761 | 133<br>1.36E-9 | 133<br>2.28E-7 | 133<br>4.37E-8 | 118<br>4.97E-4 |
| *svh-2* | volume | 69<br>0.174 | 122<br>0.00169 | 122<br>0.319 | 121<br>0.395 | 120<br>0.1 |
| *svh-2* | width | 69<br>0.174 | 122<br>4.15E-6 | 122<br>1.10E-10 | 121<br>9.11E-9 | 120<br>7.85E-7 |
| *svh-2* | length | 69<br>0.0015 | 122<br>5.65E-12 | 122<br>5.77E-14 | 121<br>9.31E-10 | 120<br>3.86E-9 |
| *mlk-1* | volume | 67<br>0.989 | 128<br>0.558 | 128<br>0.00864 | 128<br>0.0748 | 126<br>8.68E-10 |
| *mlk-1* | width | 67<br>0.0316 | 128<br>6.60E-15 | 128<br>1.13E-22 | 128<br>1.38E-23 | 126<br>4.55E-24 |
| *mlk-1* | length | 67<br>2.92E-6 | 128<br>2.01E-11 | 128<br>4.21E-16 | 128<br>9.71E-25 | 126<br>3.20E-30 |
| *daf-16* | volume | 63<br>0.349 | 86<br>1.55E-10 | 86<br>1.47E-16 | 86<br>2.27E-18 | 86<br>5.86E-18 |
| *daf-16* | width | 63<br>0.074 | 86<br>1.57E-4 | 86<br>6.84E-4 | 86<br>1.00E-7 | 86<br>2.87E-9 |
| *daf-16* | length | 63<br>0.555 | 86<br>1.18E-10 | 86<br>9.55E-11 | 86<br>1.63E-10 | 86<br>1.61E-11 |

**Figure 5h / Supplemental Figure 6: Epi RAGA-1 AID; *vhp-1(sa366)***

EV reference n (Hatch–M4): 310, 426, 426, 413, 392. Stage cells show n for the RNAi condition and exact P versus the matched EV group.

| RNAi | Trait | Hatch | M1 | M2 | M3 | M4 |
| --- | --- | --- | --- | --- | --- | --- |
| *tir-1* | volume | 93 | 150 | 147 | 146 | 144 |
|  |  | 0.425 | 2.68E-18 | 3.44E-36 | 7.90E-48 | 6.51E-35 |
| *tir-1* | width | 93 | 150 | 147 | 146 | 144 |
|  |  | 3.32E-9 | 0.172 | 1.20E-9 | 1.62E-17 | 9.39E-22 |
| *tir-1* | length | 93 | 150 | 147 | 146 | 144 |
|  |  | 0.287 | 0.00711 | 0.713 | 0.611 | 2.68E-5 |
| *nsy-1* | volume | 79 | 150 | 150 | 150 | 149 |
|  |  | 0.186 | 3.32E-20 | 3.52E-44 | 1.84E-43 | 1.59E-16 |
| *nsy-1* | width | 79 | 150 | 150 | 150 | 149 |
|  |  | 0.0578 | 7.08E-12 | 1.19E-21 | 2.62E-23 | 1.72E-20 |
| *nsy-1* | length | 79 | 150 | 150 | 150 | 149 |
|  |  | 0.544 | 5.22E-10 | 9.00E-4 | 1.24E-6 | 9.65E-30 |
| *svh-2* | volume | 93 | 124 | 124 | 122 | 121 |
|  |  | 0.0921 | 8.71E-7 | 2.37E-7 | 2.51E-9 | 2.59E-6 |
| *svh-2* | width | 93 | 124 | 124 | 122 | 121 |
|  |  | 0.0216 | 0.148 | 0.398 | 0.147 | 0.00314 |
| *svh-2* | length | 93 | 124 | 124 | 122 | 121 |
|  |  | 9.47E-4 | 3.29E-4 | 6.45E-7 | 4.23E-5 | 0.0461 |
| *mlk-1* | volume | 110 | 143 | 142 | 138 | 136 |
|  |  | 0.0333 | 2.79E-6 | 4.69E-7 | 4.08E-7 | 6.95E-4 |
| *mlk-1* | width | 110 | 143 | 142 | 138 | 136 |
|  |  | 0.172 | 0.407 | 0.705 | 0.214 | 0.0406 |
| *mlk-1* | length | 110 | 143 | 142 | 138 | 136 |
|  |  | 1.00E-4 | 3.66E-5 | 1.15E-6 | 0.0223 | 0.38 |
| *daf-16* | volume | 92 | 103 | 103 | 98 | 98 |
|  |  | 0.0436 | 0.885 | 4.84E-16 | 1.08E-13 | 1.20E-9 |
| *daf-16* | width | 92 | 103 | 103 | 98 | 98 |
|  |  | 0.0043 | 4.70E-6 | 5.10E-4 | 3.51E-4 | 1.72E-5 |
| *daf-16* | length | 92 | 103 | 103 | 98 | 98 |
|  |  | 0.162 | 7.10E-4 | 0.63 | 0.106 | 0.398 |

Deviation P values are two-sided Wilcoxon rank-sum tests comparing per-animal deviations with the same-genotype/background EV group at the indicated ecdysis. No 3×IQR filter was applied to the scaling analyses.

**Supplementary Table S8. Sample sizes and statistical results for Figure 7b-d.** For b-c, values were filtered using the 3×IQR criterion on log-transformed values. FC = median(VHP-1 AID)/median(no VHP-1 AID).

**Figure 7b-c**

| RAGA-1 background | Duration<br>N; n | Duration<br>FC; P | Growth<br>N; n | Growth<br>FC; P |
| --- | --- | --- | --- | --- |
| No RAGA-1 depletion | 4/4; 143/133 | 1.140; 5.21E-46 | 4/4; 143/135 | 0.846; 5.33E-47 |
| Epidermal RAGA-1 AID | 7/4; 322/124 | 1.163; 2.21E-53 | 7/4; 323/125 | 0.831; 9.32E-57 |

Two-sided Wilcoxon rank-sum tests. Interaction ratio: duration 1.057 (P = 5.91E-8); growth 0.945 (P = 2.51E-7). N/n values are shown for no VHP-1 AID / VHP-1 AID.

**Figure 7d: VHP-1 AID versus matched control within each RAGA-1 background**

| Background | Trait | Hatch | M1 | M2 | M3 | M4 |
| --- | --- | --- | --- | --- | --- | --- |
| No RAGA-1 depletion | volume | 98/92<br>0.4 | 145/138<br>7.12E-5 | 145/138<br>1.42E-8 | 145/138<br>9.79E-14 | 144/138<br>4.31E-14 |
| No RAGA-1 depletion | width | 98/92<br>1 | 145/138<br>0.548 | 145/138<br>0.00165 | 145/138<br>2.60E-8 | 144/138<br>9.31E-8 |
| No RAGA-1 depletion | length | 98/92<br>0.314 | 145/138<br>0.63 | 145/138<br>1.26E-8 | 145/138<br>1.47E-20 | 144/138<br>1.13E-16 |
| Epidermal RAGA-1 AID | volume | 237/81<br>0.436 | 340/145<br>2.11E-4 | 340/142<br>1.46E-5 | 339/136<br>1.48E-10 | 327/126<br>1.37E-9 |
| Epidermal RAGA-1 AID | width | 237/81<br>0.668 | 340/145<br>0.0192 | 340/142<br>5.08E-12 | 339/136<br>3.94E-30 | 327/126<br>5.26E-22 |
| Epidermal RAGA-1 AID | length | 237/81<br>0.965 | 340/145<br>0.745 | 340/142<br>0.127 | 339/136<br>2.20E-33 | 327/126<br>5.77E-32 |

Stage cells show n (no VHP-1 AID / VHP-1 AID) and exact two-sided Wilcoxon P.
